# Distinct mechanoreceptor *pezo-1* isoforms modulate food intake in the nematode *Caenorhabditis elegans*

**DOI:** 10.1101/2021.05.24.445504

**Authors:** K Hughes, A Shah, X Bai, J Adams, R Bauer, J Jackson, E Harris, A Ficca, P Freebairn, S Mohammed, EM Fernández, C Bainbridge, MA Brocco, W Stein, AG Vidal-Gadea

## Abstract

Two PIEZO mechanosensitive cation channels, PIEZO1 and PIEZO2, have been identified in mammals, where they are involved in numerous sensory processes. While structurally similar, PIEZO channels are expressed in distinct tissues and exhibit unique properties. How different PIEZOs transduce force, how their transduction mechanism varies, and how their unique properties match the functional needs of the tissues they are expressed in remain all-important unanswered questions. The nematode *Caenorhabditis elegans* has a single PIEZO ortholog (*pezo-1*) predicted to have twelve isoforms. These isoforms share many transmembrane domains but differ in those that distinguish PIEZO1 and PIEZO2 in mammals. We used transcriptional and translational reporters to show that putative promoter sequences immediately upstream of the start codon of long *pezo-1* isoforms predominantly drive GFP expression in mesodermally derived tissues (such as muscle and glands). In contrast, sequences upstream of shorter *pezo-1* isoforms resulted in GFP expression primarily in neurons. Putative promoters upstream of different isoforms drove GFP expression in different cells of the same organs of the digestive system. The observed unique pattern of complementary expression suggests that different isoforms could possess distinct functions within these organs. We used mutant analysis to show that pharyngeal muscles and glands require long *pezo-1* isoforms to respond appropriately to the presence of food. The number of *pezo-1* isoforms in *C. elegans*, their putative differential pattern of expression, and roles in experimentally tractable processes make this an attractive system to investigate the molecular basis for functional differences between members of the PIEZO family of mechanoreceptors.

## Introduction

The ability to accurately sense and respond to mechanical forces is of utmost importance for a wide range of biological processes important for survival. Mechanoreceptor proteins are instrumental in sensing mechanical forces, and they are the most versatile class of sensory transductors. Mechanoreceptors play crucial roles in many processes ranging from gating predator-prey interaction (e.g. tail flip by crayfish, or C-start escape by fish) (Currie, 1991; Katoh et al., 2013) to monitoring limb position (Woo et al., 2015a) and internal organ pressure (e.g. crop volume in flies) (Wang et al., 2020). The discovery of the mammalian mechanotransduction channel proteins (PIEZO) in 2010 not only fueled a rush of mechanotransduction research (Coste et al., 2010), but recently was also acknowledged by the 2021 Nobel Prize in Physiology or Medicine (Julius and Patapoutian, 2021). PIEZO receptors have been implicated in sensory transduction across taxa. In mammals, PIEZOs mediate touch sensation (through Merkel cells and the nerve cells that innervate them) (Vásquez et al., 2014; Woo et al., 2014), muscle length sensation (through muscle spindles) (Woo et al., 2015b), micturition in the bladder (Marshall et al., 2020), pain sensation and more (Chesler et al. 2016; Zhang et al., 2019).

The two mammalian PIEZO channels (PIEZO1 and PIEZO2) are large (840 kDa) homotrimeric proteins assembled from three ~280kDa monomers. Each monomer contributes to a central nonselective cation channel and has a peripheral domain that crosses the membrane dozens of times (Soattin et al., 2016). While similar in structure, the two mammalian PIEZO channels have distinct mechanically-activated currents and are differentially expressed. PIEZO1 is expressed in lungs, bladder, pancreas, blood cells, vascular tissue and skin, while PIEZO2 is highly expressed in sensory dorsal root ganglia and other neural tissues (Coste et al., 2010). The exact mechanisms for force transduction in PIEZO receptors are not completely understood. Two different non-exclusive models propose force transduction by PIEZOs: the force-from-lipid and the force-from-filament paradigms (Cox et al., 2017). In the force-from-lipid paradigm the channel is activated by deformations on the associated lipid membrane that surrounds the peripheral propeller part of the channel. Work with PIEZO1 receptors endogenously expressed in different cell types (Romero et al., 2019) and overexpressed in HEK293 cells (Ridone et al., 2020) suggest that these channels indeed operate according to the force-from-lipid paradigm (Syeda et al., 2016). Although PIEZO1 and PIEZO2 channel functions are modulated by the plasma membrane (Ridone et al., 2020; Romero et al., 2019), PIEZO2 appears to require an intact cytoskeleton for normal function, lending support to the force-from-filament paradigm. However, it remains unresolved if PIEZO1 requires such cytoskeletal interaction (Eijkelkamp et al., 2013; Romero et al., 2019; Wang et al., 2020). Functional differences between PIEZO1 and PIEZO2 appear to be the result of differences in the sequence of their beam (i.e., the long helix connecting the blades with the pore domain; Romero et al., 2019). In addition, PIEZO2 is differentially spliced to produce at least 16 isoforms with unique properties that are themselves expressed in specific tissues (Szczot et al., 2017). It is not presently known if differences between the transmembrane domains of PIEZO channels contribute to their function, and whether these differences lead to distinct patterns of expression.

The nematode *C. elegans* has a single ortholog of PIEZO receptors (*pezo-1*) recently shown to be mechanosensitive (Millet et al., 2021). *pezo-1* has twelve predicted isoforms (a-l, Figure 1A). Like PIEZO receptors (Figure 1B), PEZO-1 has a highly conserved pore domain and a blade domain comprising 36 (predicted) transmembrane helices (Want et al., 2019). PEZO-1 is expressed throughout the body of *C. elegans*, including the somatic sheath cells and spermathecal cells where it plays an important role in ovulation (Bai et al., 2020). All 12 *pezo-1* isoforms share the conserved pore domain but differ in the number of transmembrane segments associated with the blade component of the channels (Figure 1C). It is not presently known if *pezo-1* isoforms are differentially expressed, or the extent to which they might display functional specializations resembling those seen in the mammalian PIEZO1 and PIEZO2. Using transcriptional reporter strains we found that GFP expression driven by putative promoter regions upstream of short *pezo-1* isoform start codon occurred primarily in the nervous system, while GFP expression driven by putative promoters upstream of the start codon of long *pezo-1* isoforms largely took place in mesodermal tissues. Differential expression was also apparent in tissues of the digestive system including the pharyngeal gland cells, where RNAi-knock down of long *pezo-1* isoform expression resulted in loss of food-dependent gland activation. We propose *C. elegans* as an amenable *in vivo* system for the physiological and behavioral study of one of the most complex mechanoreceptors known. Insights about PEZO-1 isoform function will inform our understanding of the way PIEZO1 and PIEZO2 operate in mammals.

**Figure 1.**
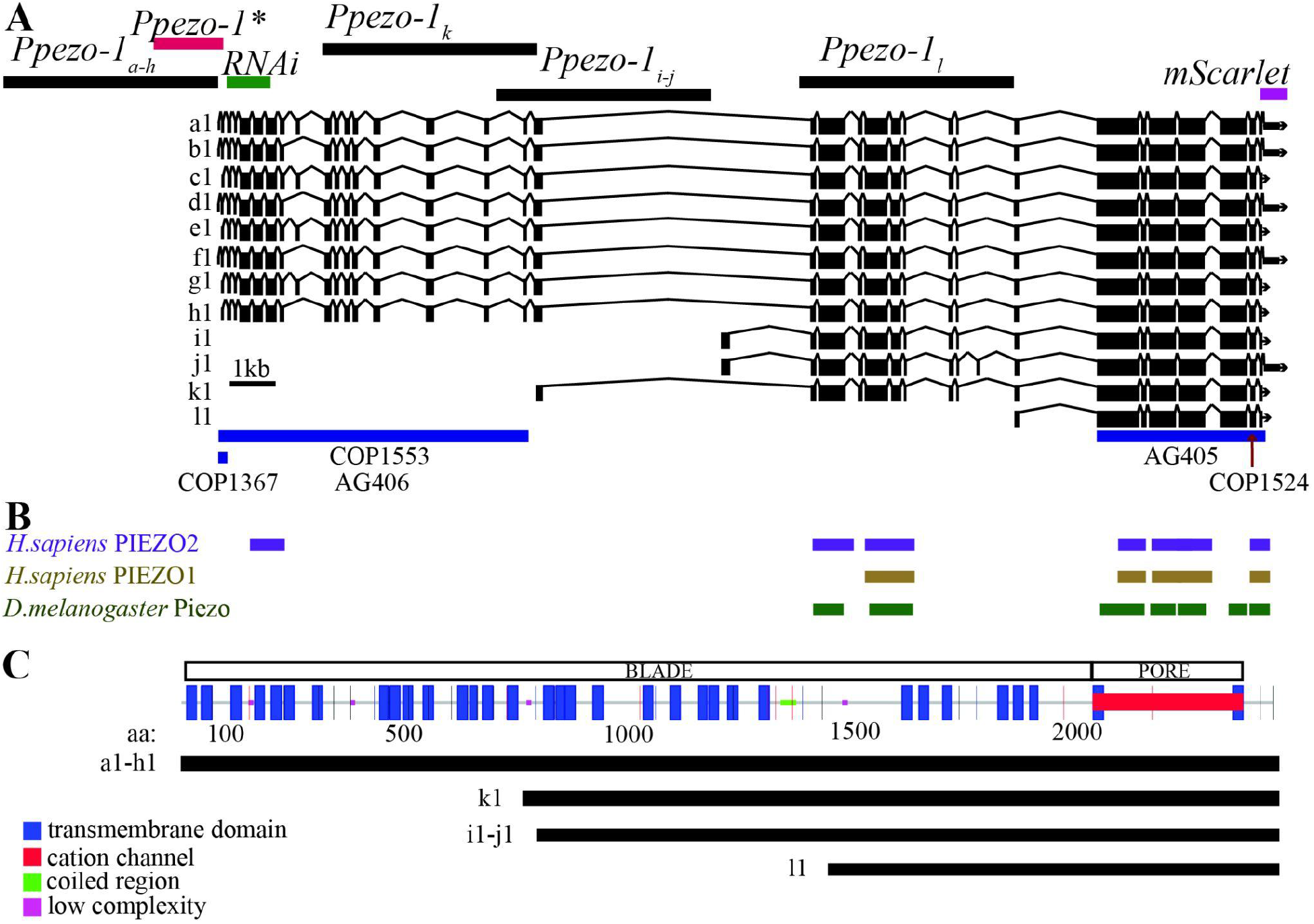
Genetic map of *C. elegans’ pezo-1*. **A)** Wormbase (Harris et al., 2020) diagram of *pezo-1* showing twelve predicted isoforms (a-l). Exons are shown in black boxes separated by introns (black lines). The 5kb sequences used to drive expression of GFP are shown targeting a-h isoforms (top left black bar), i-j, k, and l isoforms (top black bars). Also highlighted is the abbreviated promoter sequence (*Ppezo-1\**) used to drive GCaMP6s expression in the pharyngeal gland cells (red bar), as well as the coding sequence targeted by RNA interference (green bar). Under the diagram of *pezo-1* we have indicated the location of each CRISPR mutation in blue bars (COP1367, COP1553, AG406, AG405) and red arrows (COP1524). **B)** Protein similarity regions between the worm PEZO-1 and the human PIEZO1 and PIEZO2 (PIEZO1: score=100, E-val=4e-17, %ID=32.7; PIEZO2: score=82, E-val=1e-11, %ID=58.33), and the fruit fly Piezo (score=666, E-val=0.0, %ID=27.8) obtained from WormBase through ENSEMBL, and BLASTX (Yates et al., 2020; Harris et al., 2020). The bars are shown in alignment to the gene sequence in A. **C)** Functional protein domain analysis obtained through SMART and GenomeNet Motif showing the predicted location of the cation channel (red) and the transmembrane domains that constitute the propeller blade domains of the channel (Kanehisha et al., 2020; Schultz et al., 2000). The different isoform groups are aligned to show the predicted transmembrane domains present in each group (black lines).

## Results

The WormBase annotated genome browser identifies up to twelve predicted *pezo-1* isoforms (Figure 1A) (Harris et al., 2020). Protein comparisons of PEZO-1 with human and flies (using Ensembl and aligned nucleotide to protein with BLASTX) returned the human PIEZO1 and PIEZO2, and the *Drosophila* Piezo proteins (Figure 1B) (Harris et al., 2020; Yates et al., 2020). All isoforms share the C-terminus which encodes the conserved pore domain and differ mostly in the number of transmembrane segments within the blade domain of PIEZOs channels (Figure 1C) (Kamajaya et al., 2014). In *C. elegans*, the different *pezo-1* isoforms can be divided into long and short isoform groups. The long isoform group comprises the eight longest isoforms (a-h) with over thirty transmembrane segments. The four remaining shorter isoforms (i-l) have between eight and twenty transmembrane segments and differ in their predicted translational start sites.

### Promoter fusion constructs targeting different *pezo-1* isoforms have complementary expressions in gastrointestinal tissues

GFP expression driven by promoters targeting different *pezo-1* isoforms was detected in tissues associated with the digestive and reproductive systems. However, while expression was largely colocalized to the same organ systems, GFP expression patterns within these organs were distinct in strains targeting different isoforms. Below we provide a description of the tissues we identified as activating the promoter and expressing GFP which we imaged using a Leica SP8 confocal microscope.

### Transcriptional reporters targeting long *pezo-1*_a-h_ isoforms

Using a promoter immediately upstream of the start codon of the eight longest *pezo-1* isoforms we drove expression of GFP in several mesodermal tissues (Figure 2), including the body wall and pharyngeal muscles, pharyngeal glands and valve, and uterine and defecation tissues. Our findings are consistent with those described in a recent preprint by Millet et al. (2021). Based on their unique morphology and location, we identified GFP expression in pharyngeal muscles (pm3-8), pharyngeal glands (dg1, vg1s, vg2s), pharyngeal-intestinal valve (vpi) cells, AMsh and AMso glial cells (Figure 2B-D). We also tentatively identified pharyngeal neurons M1 and I4 (Figure 2E). Medially, we saw labelling of the HSN neuron, the vulval muscles, uterine muscles, and intestine (Figure 2G). Posteriorly, we saw labelling in cells involved in defecation (Figure 2H). Based on their morphology and position, we tentatively identified these as rectD, K, F, B, U, and Y (N=21) (Figure 2H). In addition to GFP-based transcriptional reporters, we used the CRISPR/Cas9 generated AG467 (integrated) strain which used mScarlet to tag the C terminus of PEZO-1 proteins. Imaging of AG467 (N=6) showed PEZO-1 proteins localized to the vicinity of the pharyngeal grinder, to the quadrants of the pharyngeal bulb occupied by the gland cells, and to the tip of the nose of the worm where the amphid sheath is located (Figure 2I).

**Figure 2.**
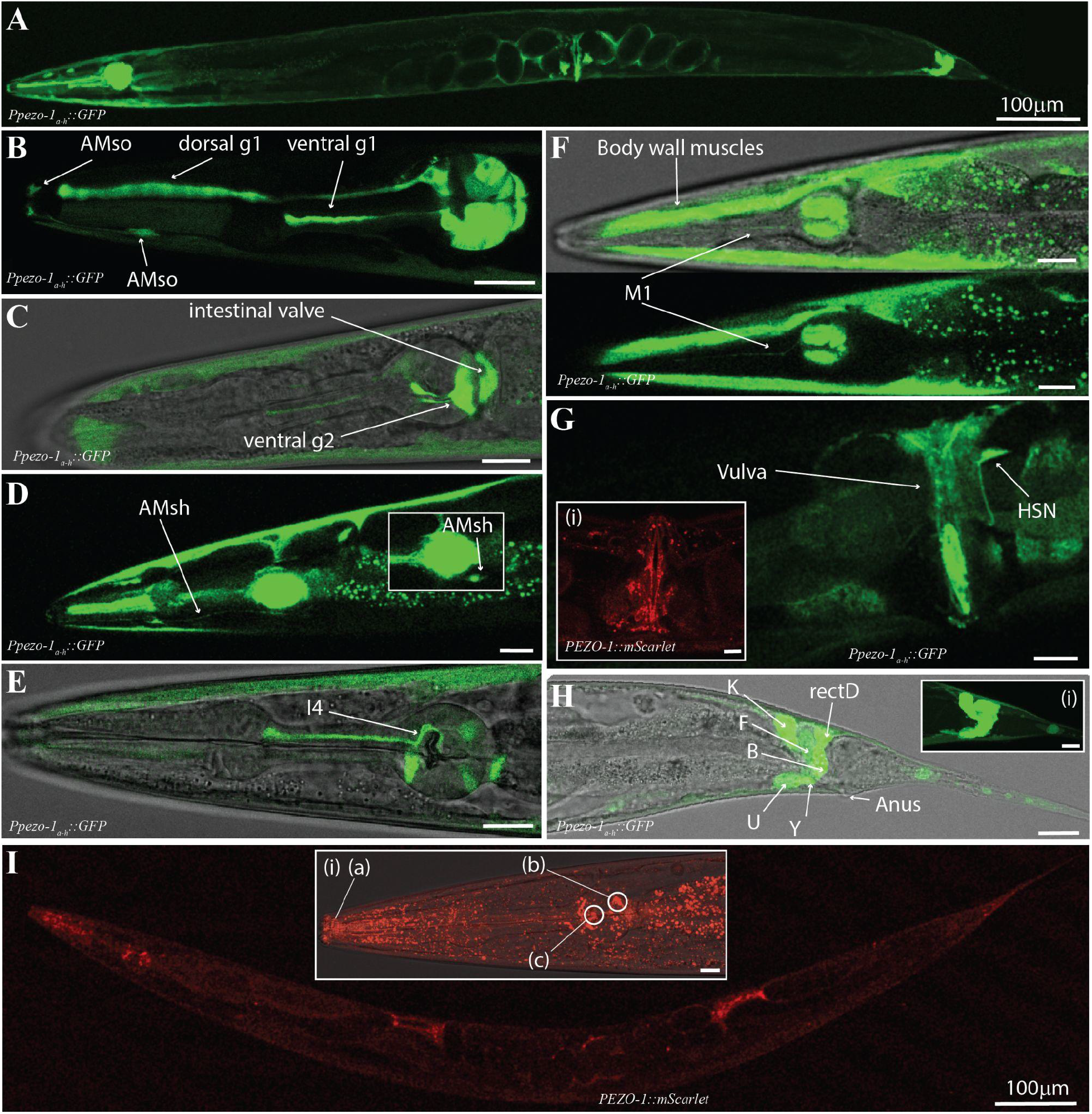
Enhancer regions upstream of *pezo-1_a-h_* isoforms start codon drove GFP expression in gonad, digestive tract, and associated tissues. **A)** A 5kb promoter upstream of the *pezo-1* start codon was used to drive expression of GFP targeting long isoform (a-h) expression. **B)** GFP was expressed in pharyngeal gland cells (dorsal and ventral g1), and the AMso glia. **C)** GFP was also present in ventral pharyngeal gland cells (vg2), the intestinal valve, and the AMsh glia **(D)**. **E)** Cells with anatomy and location consistent with the pharyngeal neuron I4 were labelled. **F)** The pharyngeal neuron M1, body wall musculature, and intestine also expressed GFP. Pharyngeal muscles including the isthmus and terminal bulb (pm3, and pm5-8) were labelled in most preps. **G)** Midbody we found GFP expression in the uterus and vulva, as well as in the HSN neuron. The AG467 translational reporter strain tagging the C-terminus of *pezo-1* proteins (PEZO-1∷mScarlet) shows PEZO-1 protein accumulation in the vulva and vulval muscles **(i)**. **H)** GFP expression was detected in several cells associated with defecation, including cells with anatomy and location consistent with K, F, B, U, Y, and rectD. **I)** The translational reporter strain targeting the C domain (shared between all isoforms) showed expression consistent with our transcriptional reporters, with obvious protein accumulation in regions consistent with the glial terminal processes **(a)**, pharyngeal glands **(b)**, and pharyngeal grinder **(c)**. Animals are presented in sagittal views (dorsal up, anterior left) with the exception of F (partial coronal projection). All scale bars are 10μm unless otherwise indicated. Observations reported based on N=21 extrachromosomal (transcriptional reporter) and N=6 integrated (translational reporter) animals. All images obtained using a Leica SP8 white light laser confocal microscope.

### Transcriptional reporters targeting short *pezo-1*_i-j_ isoforms

We used a putative promoter ~5kb sequence immediately upstream of the start codon of *pezo-1* isoforms i and j (Figure 1A). This resulted in a pattern of GFP fluorescence distinct from the one obtained above. GFP expression was mainly restricted to pharyngeal muscles and sensory neurons (N=9, Figure 3A-F). In the pharynx we saw GFP labeling of the pm3 muscle cells, the mc1 marginal cells in the corpus, a structure responsible for generating suction forces during the ingestion of suspended bacteria (Figure 3B) (Raizen et al., 2012), and in a neuron consistent in location and morphology with I4 (Figure 3C). A ventral cell located posterior to the terminal bulb and extending a process dorso-anteriorly toward the pharyngeal-intestinal valve (consistent with the excretory gland) was also labelled (asterisk in Figure 3F). Based on their location and characteristic extensive branching, we identified the anterior and posterior mechanoreceptors FLP and PVD (respectively) as also expressing GFP (Figure 3B, E).

**Figure 3.**
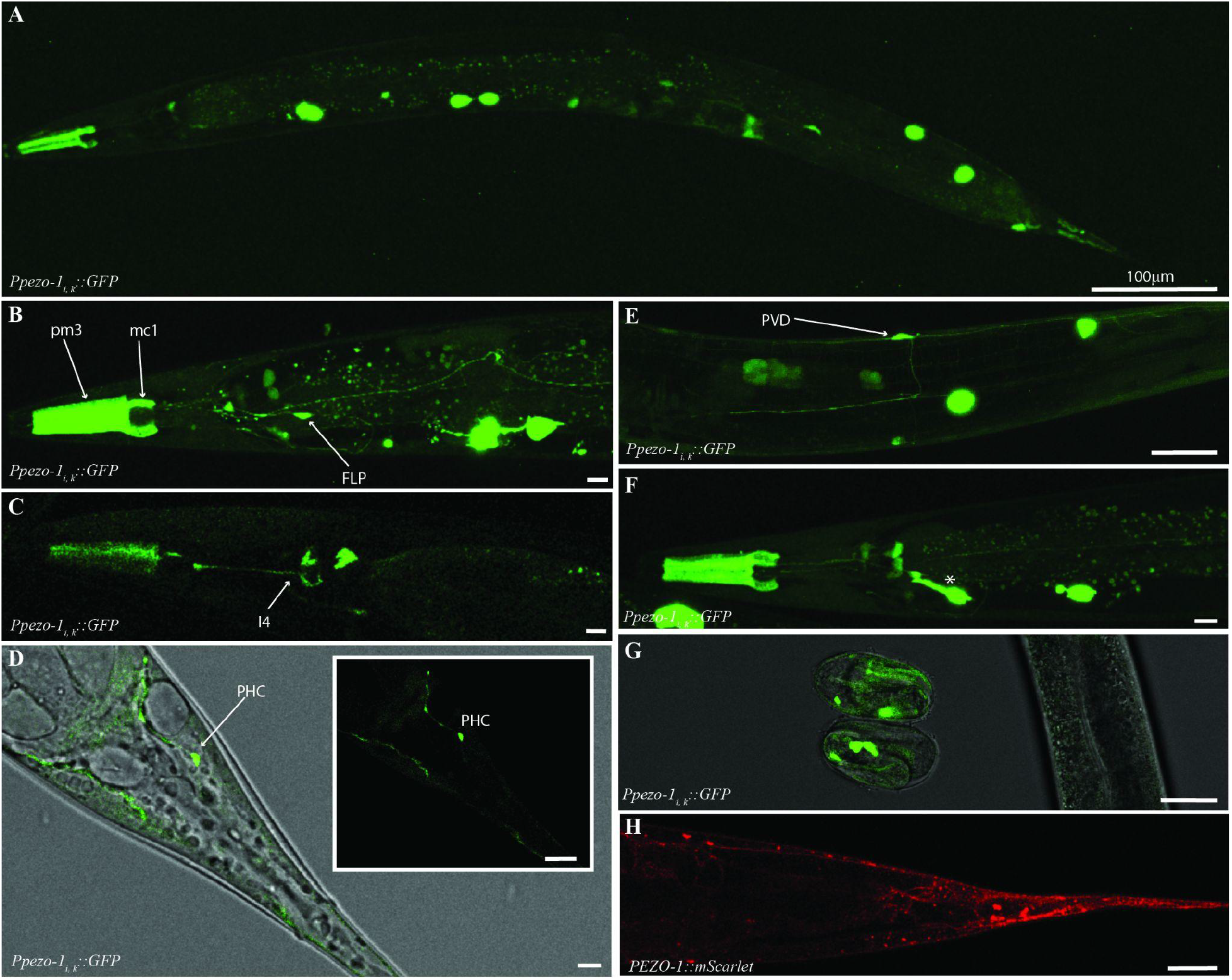
Enhancer regions upstream of *pezo-1_ij_* isoforms start codon drove GFP expression in the pharynx, and several sensory neurons. **A)** 4.9kb promoter spanning the largest intron in the *pezo-1* gene was used to drive GFP expression of the shorter *pezo-1* isoforms (using coelomocyte∷GFP as co-injection marker). **B)** GFP expression was present in pharyngeal muscles 3 (pm3), the marginal cells 1 (mc1) of the procorpus, and the anterior mechanosensitive neuron PVD. **C)** Expression was seen in a pharyngeal neuron consistent with I3. **D)** Expression was also detected in PHC, and the mechanosensitive neurons PVD **(E)**. **F)** An anterior ventral cell resembling the excretory gland was also labelled (*). **G)** Unlike GFP expression driven by promoters targeting long isoform, fluorescence signals were evident in late embryos. **H)** These expression patterns were confirmed by the PEZO-1∷mScarlet (red) strain using magenta to tag the C-terminal end of all PEZO-1 proteins. All scale bars are 10μm unless otherwise indicated. Animals are presented in sagittal views (dorsal up, anterior left). Observations reported based on N=9 extrachromosomal (transcriptional reporter) and N=6 integrated (translational reporter) animals. All images obtained using a Leica SP8 white light laser confocal microscope.

Posteriorly we tentatively identified a phasmid neuron consistent with PHC’s morphology and location, as well as other cells associated with the uterus and anus, as expressing GFP (Figure 3D). Unlike the other transcriptional constructs, GFP expression in these animals was visible as early as prezle-stage embryos (Figure 3G). As with constructs targeting longer isoforms, we observed similar patterns between promoter-driven GFP expression, and mScarlet-tagged PEZO-1 localization (Figure 3H).

### Transcriptional reporters targeting additional *pezo-1* isoform expression

We used a promoter immediately upstream of the start codon of *pezo-1* isoform k to drive GFP expression. We saw labelling of pharyngeal isthmus muscles, pharyngeal glands, and cells with morphology and position consistent with arcade cells (Figure 4A). We also saw GFP labeling in ventral neurons and in body wall muscles, where GFP colocalized with a *Pmyo-3∷mCherry* body wall co-injection marker (N=6, Figure 4B).

**Figure 4.**
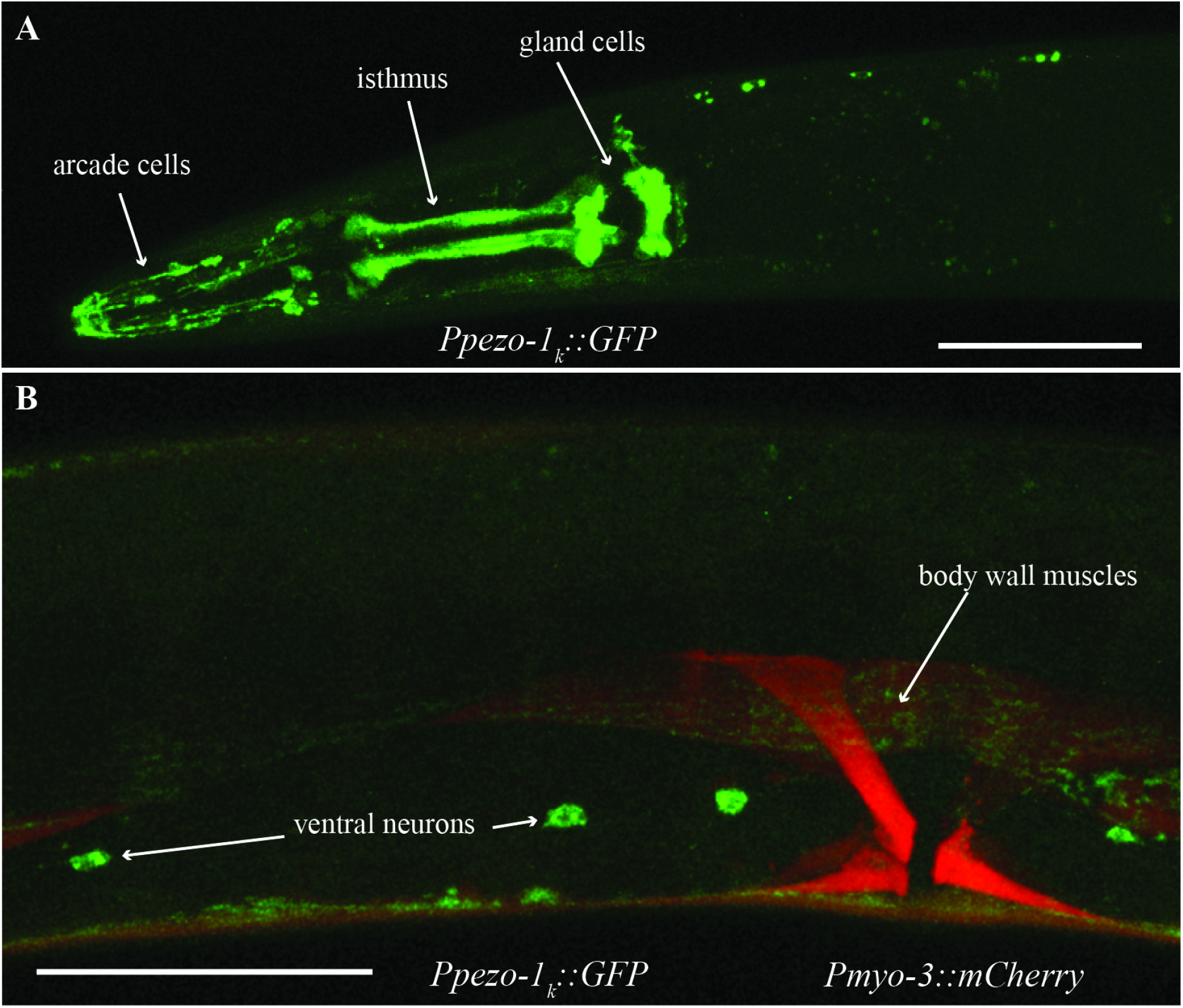
Enhancer region upstream of *pezo-1_k_* isoform start codon drove GFP expression in pharyngeal and neural tissues. **A)** GFP was observed in several pharyngeal tissues including the pharyngeal glands, pharyngeal isthmus, arcade cells, the pharyngeal valve, and AMso cells. **B)** GFP was also expressed in body wall muscles and several ventral neurons. Green is *Ppezo-1k∷GFP*, red is *Pmyo-3∷mCherry* labelling body wall and vulval musculature. Scale bar is 50μm. Observations reported based on N=6 extrachromosomal animals. All images obtained using a Leica SP8 white light laser confocal microscope.

We also used a promoter region spanning ~5kb upstream of the start codon of the shortest proposed isoform of *pezo-1*, l (Figure 1A, Supplementary Table 2); however, we were unable to detect GFP expression in these animals. This could mean that this proposed isoform does not constitute a real isoform. Alternatively, the promoter region used to drive GFP expression could have missed a necessary regulatory sequence, or lacked the appropriate cellular environment to result in activation and GFP expression (e.g. stress, age).

### Loss of *pezo-1* long isoforms results in abnormal defecation and egg-laying frequencies

After observing GFP expression in the pharynx, anal cells, and uterus, we investigated *pezo-1* necessity for processes mediated by these tissues. To this end we used a two-pronged approach. We used RNA interference (RNAi) to reduce *pezo-1* expression in tissues that are non-refractive to RNA (e.g. muscles) (Conte et al., 2015, please see note on methods regarding the *pezo-1* RNAi clone). Additionally, we used loss of function (LF) and gain of function (GOF) mutants generated through CRISPR/Cas9. While LF strains were produced by introducing large deletions, the GOF mutant was (a kind gift from Dr. Valeria Vásquez) generated by substituting an Arg 2373 for Lys (R2373K) to match similar GOF mutations in the human Piezo1 channel that result in increased cation permeability (see characterization in Millet et al., 2021).

#### Loss of long *pezo-1* isoforms increases defecation frequency

To investigate the potential involvement of *pezo-1* in defecation, we measured <u>p</u>osterior <u>Bo</u>dy <u>c</u>ontraction (pBoc) period duration in a *pezo-1* whole-gene knockout (AG405), a whole-gene gain of function mutant (COP1524), and three *pezo-1* long isoform (a-h) knockout mutants (COP1367, COP1553, and AG406). Interestingly, while mutations targeting only long *pezo-1* isoforms resulted in shorter defecation periods (i.e. increased defecation frequency), loss of function and gain of function mutations affecting both long and short isoforms had no significant effect on pBoc period (Figure 5A, One-Way ANOVA at the P<0.5 level [F=16.78, p<0.001], followed by multiple comparisons vs control group [Holm-Sidak test] at the p=0.05 level)).

**Figure 5.**
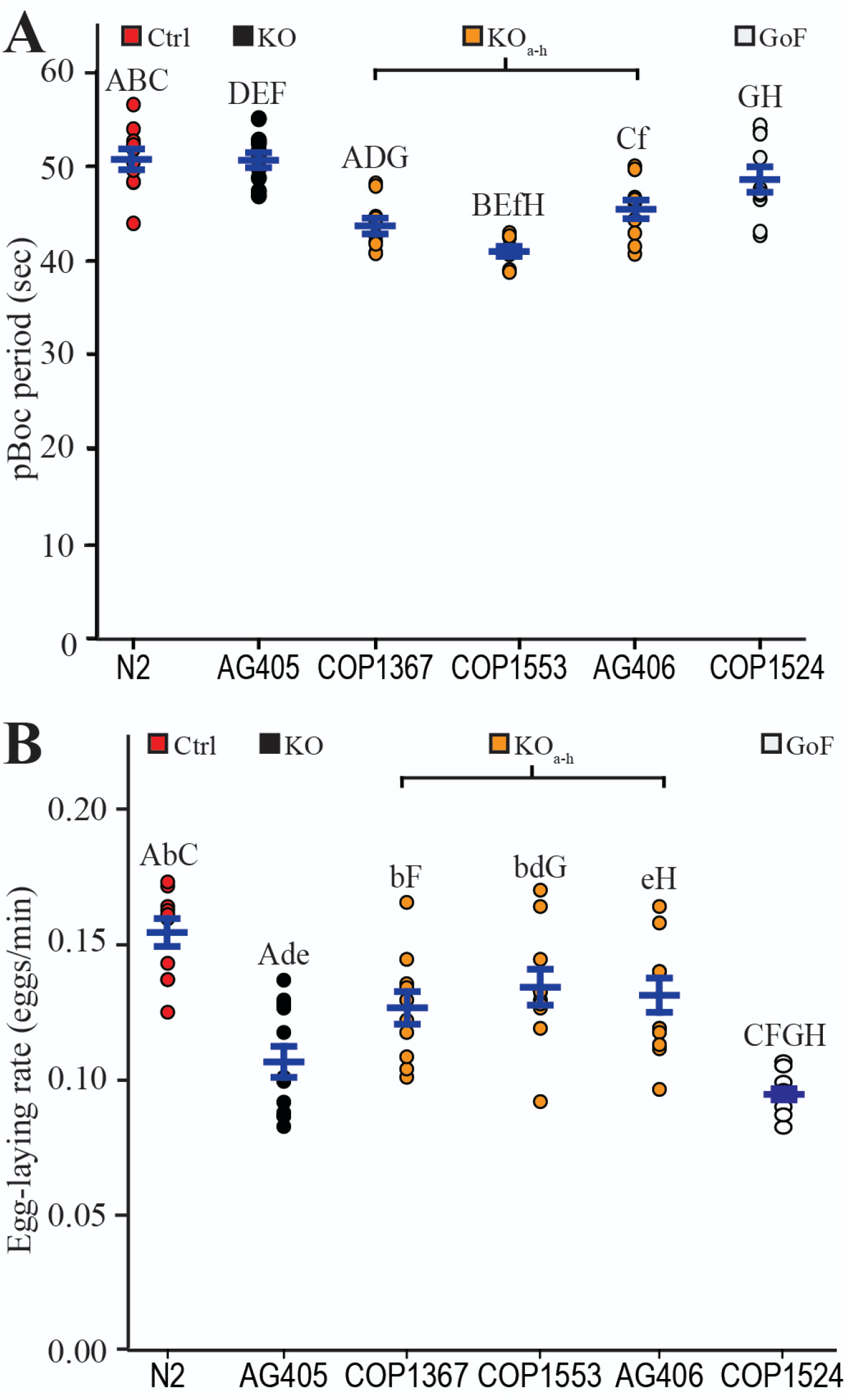
Normal defecation and egg-laying require balanced expression of *pezo-1* isoforms. **A)** Mutations resulting in loss of long *pezo-1* isoforms caused a significant decrease in posterior body contraction (pBoc) period duration. However, both loss of function and gain of function mutations affecting all *pezo-1* isoforms equally had no significant effect on pBoc period duration, N=10. **B)** Mutations affecting long *pezo-1* isoform resulted in a significant reduction in egg-laying rates for COP1367. Loss of function and gain of function mutations affecting all *pezo-1* isoforms caused an even greater reduction in egg-laying rates, N≥10. For A and B comparisons were done through One-Way ANOVA with Holm-Sidak test. Letters indicate comparisons that are of statistical significance. Values reported are means+/− SEMs. Capitalized letters indicate significance at the p<0.001 level, while lowercase letters indicate significance at the p<0.05 level. Please refer to Supplementary Table 4 for statistics.

#### Loss of *pezo-1* decreased egg-laying frequency

We next evaluated the effect of *pezo-1* mutations on egg-laying frequency. Here again we found a difference between mutations affecting all isoforms equally versus mutations affecting only long isoforms. Mutations targeting only long isoforms of *pezo-1* produced a significant decrease in egg-laying rate (One-Way ANOVA at the P<0.5 level [F=8.72, p<0.001], followed by multiple comparisons vs control group [Holm-Sidak test] at the p<0.05 level). Mutations resulting in whole-gene knockout, or a whole-gene gain of function, had a greater effect reducing egg-laying frequency than mutations affecting only long isoforms (Figure 5B, (One-Way ANOVA at the p<0.05 level [F=8.72, p<0.001], followed by pairwise comparisons between COP1367 vs AG405 and COP1524 [Holm-Sidak test] at the p<0.05 level).

### *Long isoforms* (pezo-1_a-h_) may be involved in pharyngeal pumping

Wild-type animals treated post-embryonically (L1 through adulthood) with dsRNA targeting *pezo-1*_a-h_ showed a significantly increased pharyngeal pumping frequency when compared to control treatment (L4440) worms (Figure 6A). Our behavioral experiments were carried out on standard cultivation plates and OP50 *E.coli* (food) concentrations (A600⋍0.7). However, worms are known to alter their pharyngeal pumping frequency as a function of food availability (You et al., 2006). To investigate if *pezo-1* plays a role in pharyngeal responses to changes in food density, we allowed worms to feed on OP50 *E.coli* diluted to different concentrations. We began by placing worms in a diluted bacterial lawn and transferred them into higher density lawns every 30 minutes. We found that at low food density (A600=0.69), the pharyngeal pumping rates of WT animals were not significantly different from that of animals with loss of function or gain of function mutations GCaMP6s in *pezo-1* (Figure 6B). As the bacterial lawn density increased, wild-type animals decreased their pharyngeal pumping rate (Figure 6B). However, the six *pezo-1* mutant strains displayed differing levels of inability to reduce pharyngeal pumping rates in response to increased food density (Figure 6B). A recent manuscript described a similar observation for *pezo-1* mutants when worms were fed bacteria of increasing sizes (Millet et al. 2021).

**Figure 6.**
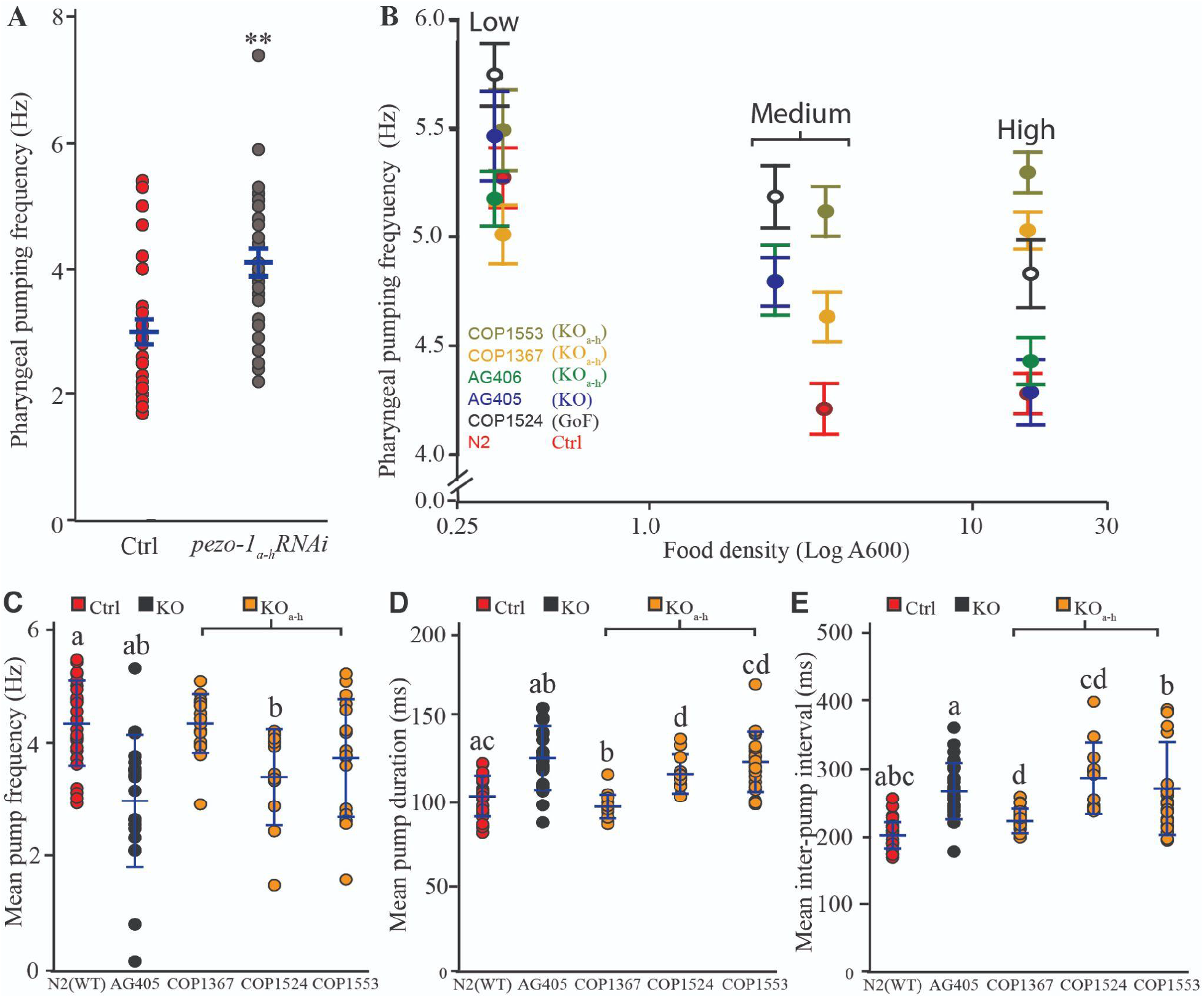
Long *pezo-1* isoforms are required for normal pharyngeal response to increases in food density. **A)** knockdown through RNAi (by feeding) of *pezo-1_a-h_* expression in a wild-type (N2) background resulted in an increase in pharyngeal pumping frequency (Mann-Whitney Rank Sum Test, N=29). **B)** Cultivation of WT (N2) *C. elegans* in bacteria lawns of increasing densities resulted in a decrease in pharyngeal pumping rate (red). However, mutations affecting long *pezo-1* isoform (a-h) resulted in worms that displayed impaired responses to increased food density (N=10). For statistical comparison we grouped the data into **Low**(A600=0.32-0.34), **Medium**(A600=2.41-3.49), and **High**(A600=14.82-15.14) density. **C)** Electropharyngeogram recordings of *pezo-1* mutants induced to perform pharyngeal pumping by immersion in 5mM serotonin solution also displayed pharyngeal pumping deficits including decreased pumping frequency, increased pump duration (**D**), and increased inter-pump intervals (**E**) (N=10-26). For C through E One-Way ANOVA on Ranks and Dunn’s on all Pairwise. Letters indicate comparisons that are of statistical significance. Values reported are means+/− SEMs. Please refer to Supplementary Table 4 for statistical comparisons between these groups.

To further characterize the pharyngeal pumping phenotype of *pezo-1* mutants we performed electropharyngeogram recording from wild-type (N2) and *pezo-1* mutants. Unlike unrestrained animals in our previous experiments (filmed while freely moving in bacterial lawns on agar plates), worms immersed in liquid media (e.g. inside our electropharyngeogram chips) do not normally engage in pharyngeal pumping unless they are chemically induced to do so by neuromodulators (Vidal-Gadea et al. 2011). We found that physically restrained (in a microfluidic chip) pharyngeal pumping by *pezo-1* mutants induced to by 10mM ectopic serotonin had no significant difference from wild-type controls in mean pump frequency, duration, or interpump intervals (Supplementary Figure 1). Recent work by Millet et al. (2021) showed that pharyngeal phenotypes of similarly-restrained *pezo-1* mutants was dependent on ectopic serotonin concentration. We therefore repeated our electropharyngeograms recordings reducing ectopic serotonin concentration to 5mM and found that now (under these low serotonin conditions) several *pezo-1* mutants displayed impaired pharyngeal pumping (Figure 6C-E). Interestingly, the phenotypes observed in these mechanically-immobilized, liquid-immersed, and ectopic serotonin-induced animals were the opposite of the phenotype observed in unrestrained, spontaneously feeding animals. Namely, *pezo-1* mutants here displayed lower (rather than higher) than wild-type pumping frequencies, and increased pump duration and inter-pump intervals.

### Pharyngeal gland activation decreases as food concentration increases

The pharyngeal glands are an essential component of food processing, and given the necessity of *pezo-1* for normal pharyngeal pumping, we hypothesized that it may also be involved in regulating pharyngeal gland activity. The role of pharyngeal glands during food consumption is not well understood owing to the experiment challenge of assessing their activity. To investigate the role of *pezo-1* in pharyngeal gland activity we constructed the AVG09 strain in which a truncated (1.6kb) putative promoter upstream of the *pezo-1* long isoforms start codon was used to drive expression in gland cells (Figure 7A). We used this strain to record calcium transients in the pharyngeal gland cells in freely crawling animals to assess pharyngeal gland activation during feeding. One limitation of this approach is that, due to the close proximity between pharyngeal gland cells, we were unable to achieve single cell resolution and therefore report the activation status of all five pharyngeal cells together.

**Figure 7.**
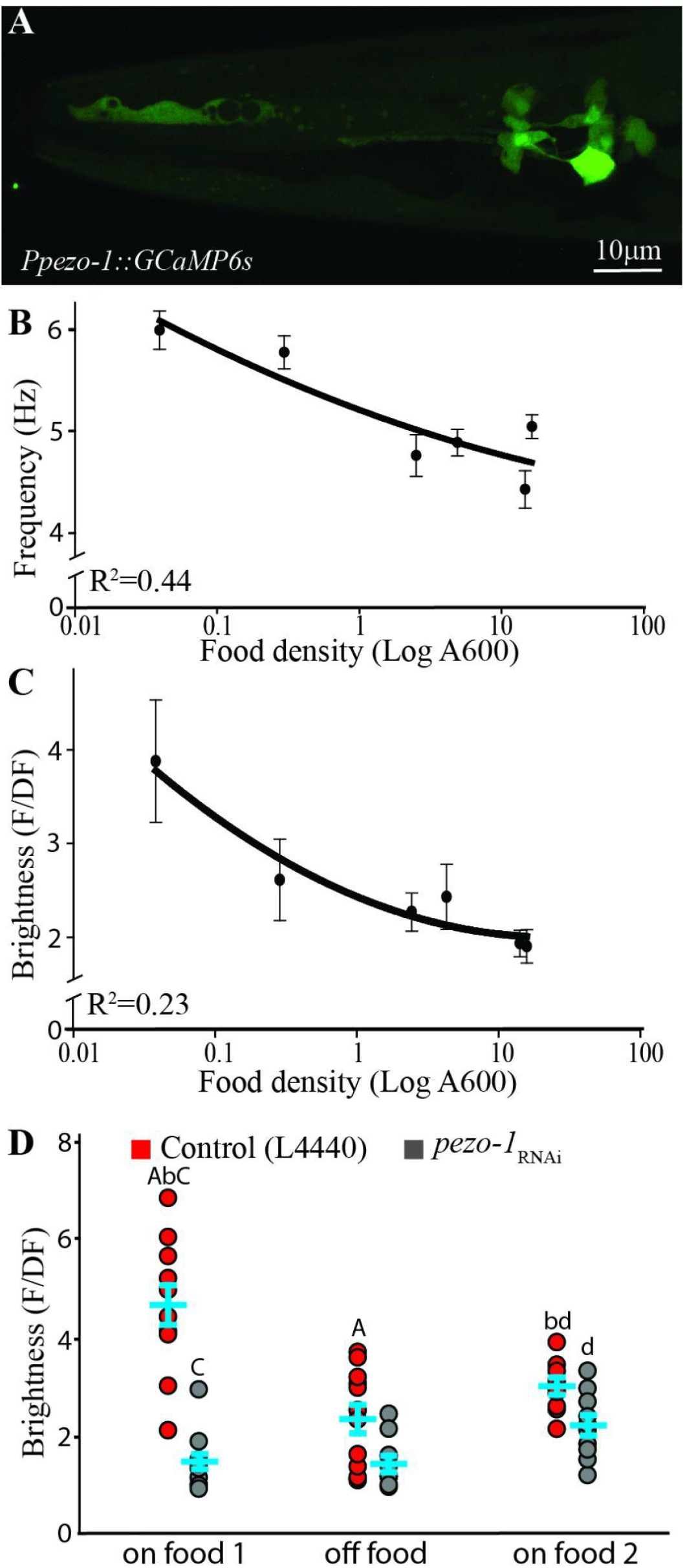
Pharyngeal gland cells respond to the presence of food in a *pezo-1* dependent manner. **A)** A short promoter 1.6kb upstream of the *pezo-1* start codon was used to drive expression of the calcium reporter GCaMP6s in pharyngeal gland cells in the strain AGV09 (expression based on N=16 animals). This allowed gland cells to be visible without obstruction by the body wall musculature (and other overlying tissues labelled with the enhancer targeting *pezo-1*_a-h_ isoforms). **B)** Increasing food density resulted in decreased pharyngeal pumping frequency, and in a decrease in gland cell brightness **(C)**, N>10. **D)** AVG09 control (L4440) and *pezo-1* RNAi-treated animals were allowed to feed for 30 minutes (*on food 1*), transferred to plates without food for 15 minutes (*off food*), and reintroduced into plates with food once again (*on food 2*). The average relative brightness of pharyngeal gland cells in control animals (red circles) decreased significantly when worms were removed from food for 15 minutes (*off food*). Knock down of *pezo-1*_a-h_ isoform expression abolished brightness changes in gland cells associated with the presence of food (grey circles). Two-Way ANOVA with Holm-Sidak pairwise comparison at p<0.05; N=10. Letters indicate comparisons that are of statistical significance. Values reported are means+/− SEMs. Capitalized letters indicate significance at the p<0.001 level, while lowercase letters indicate significance at the p<0.05 level. Please refer to Supplementary Table 4 for statistics.

We began by characterizing normal gland function. As shown above (Figure 6B) pharyngeal pump frequency is reduced in control worms when they are transferred to bacterial food of increasing density in thirty minutes intervals. Similarly, AVG09 also showed a reduction in pharyngeal pumping rate with increasing food density (Figure 7B). To test whether changes in food density also affect pharyngeal gland activity in the AVG09 strain, we cultured the worms in food of different densities and compared the fluorescence of the calcium sensor GCaMP6 in pharyngeal gland cells across different food densities. Mirroring the drop in pharyngeal pumping rate (Figure 7B), the fluorescence of pharyngeal gland cells decreased with higher food densities (Figure 7C). To ensure that the decrease in fluorescence seen in worms repeatedly transferred to plates of increasing food density was not the result of photobleaching, we repeated the experiment but this time assigned different groups of worms bacteria of different densities (rather than feeding the same worms different density bacteria sequentially as done in Figure 7C). This produced similar results as before (Supplementary Figure 2), suggesting that gland cell activation indeed decreases with increasing food density.

### *Long pezo-1*_a-h_ isoforms might be involved in food-dependent activation of pharyngeal gland cells

Our results indicate that pharyngeal pumping and pharyngeal gland activation are influenced by the density (concentration) of the bacteria they ingest. This suggests that the presence or absence of food should be reflected in pharyngeal gland activity as well. We found that in animals fed HT115 *E. coli* with the control RNAi plasmid (L4440) the brightness of the pharyngeal glands decreased significantly 15 minutes after worms were removed from food (*off food*, Figure 7D red circles). To investigate the potential role of *pezo-1* in the response of the glands to the presence of food, we repeated the experiment in *pezo-1_a-h_*-knocked down animals (fed HT115 *E. coli* with the *pezo-1* plasmid). We found that in this treatment pharyngeal gland brightness was reduced in the presence of food when compared to control animals, and did not significantly change after animals were removed from food for 15 minutes. However, Brightness increased significantly upon reintroduction to food (*on food 2*, grey circles in Figure 7D, Two-way ANOVA see Supplementary Table 4 for statistics).

Our results are consistent with normal *pezo-1* expression being important for pharyngeal pumping, defecation, and egg-laying. Furthermore, whole-gene gain and loss of function experiments suggest that loss of one class of isoforms (e.g. long) appears to be more deleterious than loss or gain of function mutations affecting all isoforms equally, hinting perhaps at an antagonistic role for different classes of isoforms in at least some of their functions.

## Discussion

*C. elegans* has a single ortholog (*pezo-1*) of the mammalian PIEZO channels which has a fair amount of conservation with human PIEZO1 and PIEZO2 as well as with the *Drosophila* Piezo channel (Figure 1). The greatest degree of conservation is for the pore domain (Kamajaya et al., 2014). Up to 12 different isoforms are predicted for *pezo-1* (Harris et al., 2020). This large number of isoforms is consistent with mammals, where PIEZO2, for example, was shown to be alternatively spliced and expressed in different tissues (Szczot et al., 2017). Alternative splicing of ion channels is also known to occur in *C. elegans* (Johnson et al., 2011). We explored the possibility that different *pezo-1* isoforms may be expressed differentially by using putative promoter regions upstream of the start codon of single or groups of isoforms to drive GFP expression. We found that the constructs targeting long *pezo-1* isoforms (a-h) displayed largely mesodermal GFP expression, similar to known PIEZO1 expression in mammals (Hennes et al., 2019). Expression included muscle, gonads, and glands (Figure 2). In contrast, constructs targeting shorter *pezo-1* isoforms resulted in GFP expression in sensory neurons (Figure 3). These results are consistent with recent findings in worms by Millet et al. (2021), and in mammals where different isoforms are expressed in different germ layers, suggesting similarly distinct functions for different isoforms in *C. elegans* (Coste et al., 2010).

Our study was not able to target individual isoforms, but rather grouped the longest eight isoforms (a-h) because of their shared start codon. The putative promoter regions for the remaining (shorter) isoforms (and their predicted start codons) are distributed across a 10kb region. For the purpose of this study, we used a putative promoter sequence immediately upstream of the i and j isoforms start codon, the k isoform start codon, and the smaller, l, isoform start codon. We designed our promoter sequences trying to maximize the region upstream of the putative isoform translational start site, while minimizing overlap with other known genes or isoforms. It is, however, possible that our constructs could have erred by either including regulatory sequences belonging to other isoforms, or by neglecting downstream regulatory regions important for the transcription of a targeted isoform. Therefore, tissues identified in this (and similarly designed) work remain putative awaiting cell-specific qPCR validation. Indeed, a CRISPR knock-in strain (AG467) which tags the fluorescent reporter mScarlet at C termini of PEZO-1 and is predicted to encompass all isoforms because of their common stop site shows broader PEZO-1 protein expression than what our combined transcriptional reporter strains identified (Bai et al., 2020). Similarly, RNA-based reports for L1-L4 larvae obtained from the CenGen database highlights additional tissues that express *pezo-1* (Supplementary Table 5, Hammarlund et al., 2018).

Consistent with the patterns of expression detected above, we found that normal expression of long *pezo-1* isoforms was required for normal defecation frequency. Specifically, a loss of long isoforms (through LF mutations) reduced defecation frequency. However, loss of all isoforms, or gain of function mutation affecting all isoforms, did not have an impact on defecation frequency. These results suggest a potential functional antagonism between long and short *pezo-1* isoforms.

Unlike defecation (where loss of *pezo-1* led to increased contraction frequencies), we found that genetic manipulations of *pezo-1* resulted in decreased egg-laying frequencies. A gain of function mutation affecting all isoforms also decreased egg-laying frequency, suggesting that *pezo-1* levels must be carefully regulated to preserve normal egg-laying function. It remains unclear if this effect is related to reports of reduced brood size for *pezo-1* knockout mutants, where brood size in mutants was tightly linked to ovulation (Bai et al., 2020).

We found that wildtype animals decreased their pharyngeal pumping frequency as the density of the food surrounding them increased (Figure 6B red, Figure 7B). The idea of animals modulating pharyngeal pumping based on the amount of food within the pharynx has been previously explored by Scholz and colleagues. These authors used a microfluidic chip and found that increasing the flow rate of liquid containing bacteria resulted in a significant decrease in pharyngeal pumping (Scholz et al. 2016). These results are consistent with our findings of potentially increased food in the pharynx leading to a decrease in its rate of pumping.

We further found that mutations targeting long isoforms of *pezo-1* expression altered pharyngeal pumping frequencies in a context-dependent manner. Loss of *pezo-1_a-h_* signaling (through mutation or RNA interference) in freely crawling worms feeding on OP50 resulted in abnormally elevated pharyngeal pumping rates when food density was high, but not when food density was low (Figure 6B). In contrast, electropharyngeograms from the same animals showed abnormally low pharyngeal pumping frequencies for *pezo-1* mutants in the presence of 5mM serotonin (Figure 6C), but no effect in the presence of 10mM serotonin (Supplementary Figure 1). Work by Millet et al. (2021) recently showed that the frequency of serotonin-induced pharyngeal pumping depends on its concentration (with higher frequencies resulting from higher serotonin concentrations). This group found a similar pharyngeal pumping phenotype for *pezo-1* mutants. Specifically, they found that mutants fed standard *E. coli* displayed greater than wild-type pharyngeal pumping frequencies, but the same animals fed large (incompletely divided) *E. coli* displayed a lower than wild-type pharyngeal pumping frequency (Millet et al., 2021).

There are several differences between pharyngeal pumping experiments that might account for the different phenotypes observed for freely moving and (ectopic) serotonin-activated *pezo-1* mutants. Considering neuromodulation, for example, worms crawling on an agar plate will slow down to feed upon entering a bacterial lawn (a response mediated by dopamine, termed *basal slowing response*). However, when worms become starved, they perform an *enhanced slowing response* which is instead mediated by serotonin (Sawin et al, 2000). We previously showed that when worms transition to liquid environments (such as the interior of a microfluidic device used for all electropharyngeogram experiments) the altered balance between dopaminergic and serotonergic modulation leads to the inhibition of many behaviors (including foraging and pharyngeal pumping, Vidal-Gadea et al 2011, 2012). Under these aquatic conditions, ectopic serotonin application is required for most electropharyngeal recordings (Lee et al 2016, Ortiz et al 2018, Millet et al. 2021). It remains unclear the extent to which ectopic serotonin-induced pharyngeal pumping in microfluidic devices resembles the endogenous modulation of unrestrained animals under normal (i.e. basal slowing) or starved (i.e. enhanced slowing) conditions. Additionally, several key differences between freely moving and ectopic serotonin-activated assays further confound their comparison. For example, worms on a plate feed on bacteria and experience the chemical and mechanical sensory inputs associated with their mastication and ingestion. In contrast, most studies of worms on microfluidic chips consist of worms feeding liquid lacking bacteria and the associated sensory (chemical and mechanical) inputs they would normally derive from them. We think this is a relevant point in the context of a mechanosensor which might be responsible for sensing food-derived mechanical forces within the pharyngeal cavity. Furthermore, in addition to mechanosensation, Millet et al (2021) report an osmolarity dependent effect for *pezo-1* mutants where increased pharyngeal frequency in *pezo-1* mutants in a microfluidic device was observed only when the pharyngeal contents were at 320 mOsm (i.e. nearly isosmolar with the worm) but not at lower osmolarities (150 and 260 mOsm). Lastly, unlike freely crawling worms, animals in a microfluidic device are physically restrained. Based on the broad expression of *pezo-1* we established in this manuscript; it is possible that mechanical immobilization within these devices would activate many *pezo-1* (and other) mechanoreceptors in the musculature and nervous system of these animals that could indirectly affect pharyngeal pumping. We therefore feel that differences in the valence of *pezo-1* phenotypes for freely crawling animals and animals in microfluidic devices are not necessarily in conflict, but might reflect differences in the way these animals are being tested. Clearly, each approach has much to contribute to our understanding of *C. elegans* gastrointestinal physiology.

While the aim of this work was not to compare or equate two different techniques, we propose a possible explanation that might reconcile unrestrained and microfluidic chip observations. High pharyngeal pumping frequencies in microfluidic devices compared to lower frequencies on bacterial lawns result from ectopic serotonin baths initiating a “post-starvation feeding program” with high frequency pumping in these devices (Avery and Horvitz 1989, Raizen et al. 2012). In this context, the absence of mechanical feedback in the pharynx (due to the lack of food) would result in a higher pharyngeal pumping rate in control animals (i.e. by mimicking low food density) and lower than control pumping rates in *pezo-1* mutants (which may be unable to detect pharyngeal contents). Conversely, for unrestrained animals in bacterial lawns (and without ectopic serotonergic modulation), absence of mechanical feedback caused by *pezo-1* mutations would result in these mutants being unable to reduce pharyngeal pumping rates in response to increased food density, leading to a higher than normal pumping rate when food density increases. Food-induced pumping modulation has been shown before. For example, worms decrease pumping frequency when transferred from bacteria of low nutritional value to bacteria of high nutritional value (You et al., 2006). Based on our combined results, we propose that in the pharynx, *pezo-1* may play a role in discerning the contents of the pharynx. If this is the case, this would be a conserved function for *pezo-1*, as recent work in *D. melanogaster* revealed that the fly ortholog of PIEZO is similarly involved in the modulation of feeding. Mutations impairing *Drosophila*’s Piezo also result in increases in pharyngeal pumping frequency (Wang et al., 2020).

The pharyngeal glands also appeared to express *pezo-1*, where it was found to be required for their normal function. The pharyngeal gland cells are thought to aid in food lubrication and digestion, and they send processes to the different pharyngeal compartments (Raharjo et al., 2011).

The role of PIEZO channels in mechanoreception constitutes a new and thriving field of inquiry. It is clear that these channels mediate an extensive array of physiological processes across taxa. In *C. elegans, pezo-1* was implicated in male mating (Brugman, 2020) and ovulation (Bai et al., 2020). The present study represents an initial investigation into the differential expression and function of *pezo-1* isoforms. In the digestive system of *C. elegans, pezo-1* may enable different organs to independently detect and respond to the presence and quality of food. PEZO-1 thus appears to reflect the mammalian PIEZOs not only in structure but also in function. *C. elegans* is thus a promising system to study how differences between different PIEZO isoforms endow these channels with the properties that tune them to the different functions and may inform studies in mammals about these properties as well.

## Methods and Materials

### Animal strains and cultivation

The following strains were used: N2 (WT), AVG09, AVG10, AVG11,, AVG22, AVG23,TU3311, COP1367, COP1553, COP1524, OH15500, AG405, AG406, AG467. Strains COP1367, COP1553, and COP1524 were a kind gift of Dr. Vásquez at the University of Tennessee Health Science Center-Memphis. For strain description refer to Supplementary Table 1. L4 larvae were picked on the day before filming. The following day, gravid hermaphrodites were tested on day one of adulthood. Unless otherwise noted, animals were cultivated on standard NGM agar plates with OP50 *E. coli* food in a room maintained within 20-22°C and between 30-37% humidity.

### Construction of strains

To determine the pattern of expression of different *pezo-1* isoforms we built several transcriptional reporter strains (Figure 1A and Supplementary Table 1). We selected the putative promoter regions immediately upstream of the start codon of targeted isoforms, and used these regions to drive expression of a green fluorescent protein (GFP) reporter. This approach has been extensively used to report the putative cellular expression patterns of target genes (Hobert, 2002; Boulin et al., 2006).

#### Transcriptional reporter strains

The putative expression pattern for the longest *pezo-1* isoforms (a-h) was assessed using a ~5kb promoter upstream of the gene start codon. There is a large intronic region (>4kb) upstream of isoforms i and j of *pezo-1*. Predicting the presence of an intronic regulatory region, we designed primers targeting isoforms i and j by amplifying the ~5kb intronic region immediately upstream of their predicted start codon (*Ppezo-1*_i-j_ in Figure 1A). We next constructed a reporter strain using the 5kb upstream of the start codon of isoform k, and similarly another ~5kb promoter sequence immediately upstream of the start codon of isoform l, the shortest predicted isoform (*Ppezo-1*_k_ and *Ppezo-1*_l_ respectively). We used standard PCR-fusion (Hobert, 2002) to merge these different putative promoter sequences to GFP to generate the reporter strains AVG10 (*Ppezo-1_a-h_∷GFP∷unc54_3’UTR*), AVG11 (*Ppezo-1_i-j_∷GFP∷unc54_3’UTR*), AVG23 (*Ppezo-1_k_∷GFP∷unc54_3’UTR*), and AVG22 (*Ppezo-1_l_:GFP∷unc54_3’UTR*). We note that the AVG strains in this work are extrachromosomal, and that for each strain we imaged multiple individuals from a single transgenic line in order to assess mosaicism or variation in array expression. Here we report the totality of labelled tissues observed across multiple animals expressing the same array (N=21, N=9, and N=6 for isoforms a-h, i-j, and k respectively). We found that individual lines expressing the same construct showed similar expression patterns.

#### Activity reporter strain

To generate the activity reporter strain targeting pharyngeal gland cells (AVG09) a truncated (1.6kb) promoter upstream of the *pezo-1* long isoforms start codon was used to drive GCaMP6s expression in gland cells (as well as uterus and anal cells) without confounding signals from overlaying body wall and pharyngeal musculature.

### Statistical analysis

All the bars report means and variation is given as SEM throughout. All statistical analysis was performed using Sigmaplot 11 (Aspire Software). Comparisons between two different treatments were performed by planned, two-tailed paired or unpaired t-tests to compare two groups that were normally distributed and had similar variance. Samples that failed the normality or equal variance tests were compared using the Mann-Whitney Rank Sum Test. Comparison between multiple parametric datasets were done using One-Way or Two-Way ANOVAs (depending on the number of treatments), and followed by Holm-Sidak all-pairwise post hoc tests. To compare non-parametric datasets we used One-Way ANOVA on Ranks. These were followed with Dunn’s post hoc tests (all Pairwise comparisons) as these accommodated different sample sizes. In all cases, p values are reported following the convention: *p<0.05 and **p<0.001. For all electrophysiological data, outliers were identified by ROUT with Q=0.5%. Please refer to Supplementary Table 4 for summary statistics for Two-Way ANOVAs, and description of tests conducted in this study. All raw measurements are available in a supplemental Excel file (Supplementary File 1).

### Bioinformatics

Protein sequence comparisons were obtained through the protein similarity tool using ENSEMBL (Yates et al., 2020) and incorporated to WormBase (Harris et al., 2020). Functional protein domains were predicted using SMART and GenomeNet Motif search tools (Kanehisa and Sato, 2020; Schultz et al., 2000). We should point out that the widely available RNAi clone targeting *pezo-1* and distributed through the Source Bioscience RNAi library was found to target an incorrect sequence. Source Bioscience provided us with a new and validated strain (see Supplementary Table 3) targeting the region of the gene consistent with isoforms *a* through *h* (N terminus, Figure 1A). This new validated clone is now available through Source Biosciences. While the dsRNA clone targets *pezo-1*_a-h_, it does not target the shorter isoforms (i through l). Gene expression levels for *pezo-1* were obtained using The *C. elegans* Neuronal Gene Expression Map & Network (CeNGEN, Supplementary Table 5, Hammarlund et al., 2018).

### Gene knockdown

RNA interference by feeding was conducted as previously described (Conte et al., 2015). Day one adult worms were allowed to lay eggs on RNAi plates for one hour, and the eggs were then grown on the same agar plates containing IPTG and seeded with bacteria containing RNAi empty control vector (L4440), or a vector targeting the gene of interest.

### Filming

We used a Flea2 camera (Point Grey, Vancouver) mounted on an Olympus SZX12 stereomicroscope to acquire sequences of (1032×776 pixels) TIFF images. Crawling animals were individually filmed at 30Hz.

### Calcium imaging

All calcium comparisons were conducted on animals filmed under identical illumination and magnification conditions (alternating strains during the same filming session). Calcium signals were recorded with the aid of an X-cite 120 PC Q light source. We used ImageJ to measure GCaMP6s signals. We measured the brightness of the pharyngeal glands, which is an indirect measurement of calcium in the cells, by drawing an area of interest around them and then obtained a background fluorescence measurement by moving said area of interest just outside the worm and measuring this area as well. We used ImageJ to obtain maximal brightness of the gland cells, the background immediately lateral to the worm near the gland cells, and by calculating and reporting the ratio of maximal brightness of the pharyngeal glands to the maximal brightness of the background.

### Confocal imaging

We used a Leica SP8 confocal microscope to acquire 40x image stacks with a 1.3 zoom. Imaged volume was 139 μm × 56.7 μm × 38.9 μm (X, Y, and Z respectively). Stacks consisted of 85 × 0.46 μm steps. Pixel sizes were 0.136 μm, 0.136 μm, and 0.457 μm for X, Y, and Z respectively. For animals with constructs targeting the *pezo-1_a-h_* isoforms we imaged 21 animals, for those targeting *pezo-1_ij_* isoforms we imaged 6 animals, and for those targeting the *pezo-1_k_* isoforms we imaged 6 animals. The number of animals imaged reflected the extent of tissue labeling and the variability of tissues tagged between animals (with strains displaying consistent and limited labelling requiring fewer samples). Confocal images were analyzed offline using Leica Application Suite X (LAS X).

### Electropharyngeograms (EPG)

10-15 mid-L4 hermaphrodites were picked onto 35 mm MYOB plates seeded with OP50 bacteria and allowed to lay eggs for six hours before all adults were removed from the plates. The synchronized F1 population was washed off from MYOB plates after 72 hours of growth at 20 °C or 48 hours at 25 °C. The collected worms were rinsed three times with M9. The worm pellets were incubated in 10 mM (or 5 mM) serotonin (InVivo Biosystems, Inc.), for at least 20 minutes. Pharyngeal pumping was recorded with the ScreenChip system (SKC101A, InVivo Biosystems, Inc.) by following a lab-optimized protocol. The automated data were acquired with NemAcquire (InVivo Biosystems, Inc.) and analyzed by NemAnalysis software (InVivo Biosystems, Inc.). All tested strains were repeated at least three times.

### Image analysis

Non-confocal images were analyzed using ImageJ.

### Behavioral experiments

#### Pharyngeal pumping

Worms were placed in agar plates on OP50 *E. coli* bacterial lawns and allowed to feed for 30 minutes prior to imaging. Animals were filmed using a FlyCap camera (Media Cybernetics) to acquire a series of TIFF images at 40X magnification and 30fps for 15 seconds. A minimum of 10 animals were recorded for each condition. Pharyngeal pumping was measured blind to treatment using ImageJ software and using the motion of the terminal pharynx grinder as the visual cue. For all pharyngeal pumping experiments (34 assays; 10-62 animals/assay; N=536), we note that sample size was not correlated with pharyngeal pumping frequency variability (Pearson Product Moment of SEM vs sample size p>0.05; and linear regression between the two variables r^2^=0.0008).

#### Defecation assay

We transferred day-1 adult hermaphrodite worms into NGM agar plates seeded with OP50 *E.coli* and allowed the worms to feed for 30 minutes before imaging. Animals (>10 per condition) were imaged at 30fps for 10 minutes. Five consecutive posterior body contractions (pBoc) were measured for each animal. The pBoc period was calculated as the average time between consecutive posterior body contractions.

#### Egg-laying assay

The rate of egg laying was assayed by placing ten day-one adult worms in a copper ring (measuring 1.4 cm in diameter) over a bacterial lawn. Worms were allowed to lay eggs for sixty minutes after which they were removed, and the number of eggs counted. We then divided the number of eggs by the number of worms, divided by the number of minutes to get eggs laid per minute. Ten assays were performed for each strain.

##### Food density assay

We centrifuged 2ml of overnight cultured OP50 *E.coli* for 2 min at 10,000 rpm and discarded the supernatant. The bacterial pellet was resuspended in 50 μl of liquid NGM and its optical density at 600nm (OD_600_) was measured using a Thermo Fisher Nanodrop. We made five serial dilutions of this stock and measured their optical density as above. A 30 μl drop of each dilution was placed on an NGM agar plate and allowed to dry for 30 minutes. Worms were transferred into the lowest OP50 density plate and allowed to feed for 30 minutes prior to imaging. Pharyngeal pumping was filmed for each worm as described above. After all (N≥10) worms were filmed they were transferred into the plate with the next higher food concentration and the process was repeated until all the animals were filmed in all food concentrations.

For measuring the activation of the pharyngeal gland cells under different food concentrations we performed the experiments as above but used a fluorescent light to film the pharyngeal gland cells (as described above). Because these animals were reimaged at increasing concentrations, we were concerned that changes in fluorescence could be indicative of fluorophore bleaching rather than the potential changes in cytoplasmic calcium we were interested in measuring. We therefore repeated this experiment but this time placing 10-15 animals in plates of each food concentration at the same time before filming them. Animals were allowed to feed for 30 minutes before filming with a fluorescent light as described above.

##### Food availability assay

For the food presence assay (10-15) animals were allowed to feed on OP50 *E. coli* lawns on NGM agar plates for 30 minutes. Their pharyngeal pumping (or pharyngeal gland fluorescence) was filmed as described above to produce the “*on food 1*” condition. The animals were then individually transferred to an NGM agar plate without food for 15 minutes. After this time their pharyngeal pumping (or pharyngeal gland fluorescence) was again filmed to produce the “off food” condition. Animals were then transferred back to a bacterial OP50 lawn where they were filmed one final time within one minute of being reintroduced to produce the “on food 2” condition.

## Author contributions

HK and VGAG: performed experiments, analyzed data, wrote manuscript. SA: performed experiments. AJ: constructed strains. XB: performed experiments, constructed strains, analyzed data. RB: performed experiments, analyzed data. JJ: performed experiments. BC: constructed strains. HE, FA, FP, MS, FEM: performed experiments. BM, SW: wrote manuscript.

## Data availability

Strains are available upon request. The authors affirm that all data necessary for confirming the conclusions of the article are present within the article, figures, and tables.

## Acknowledgements

Some strains were provided by the Caenorhabditis Genetics Center (funded by NIH Grant P40 OD010440). Funding was provided by NIH Grant 1R15AR068583-01A1, and NSF Grant 1818140 to A.G.V.-G. This work was supported, in part, by the Intramural Research Program of the National Institutes of Health, National Institute of Diabetes and Digestive and Kidney Diseases (X.B., R.B). We thank Dr. Andy Golden for feedback on the manuscript. We thank Dr. Valeria Vásquez for the worm strains.

## Supplementary Information

### Supplementary Figure Captions.

**Supplementary Figure 1.**
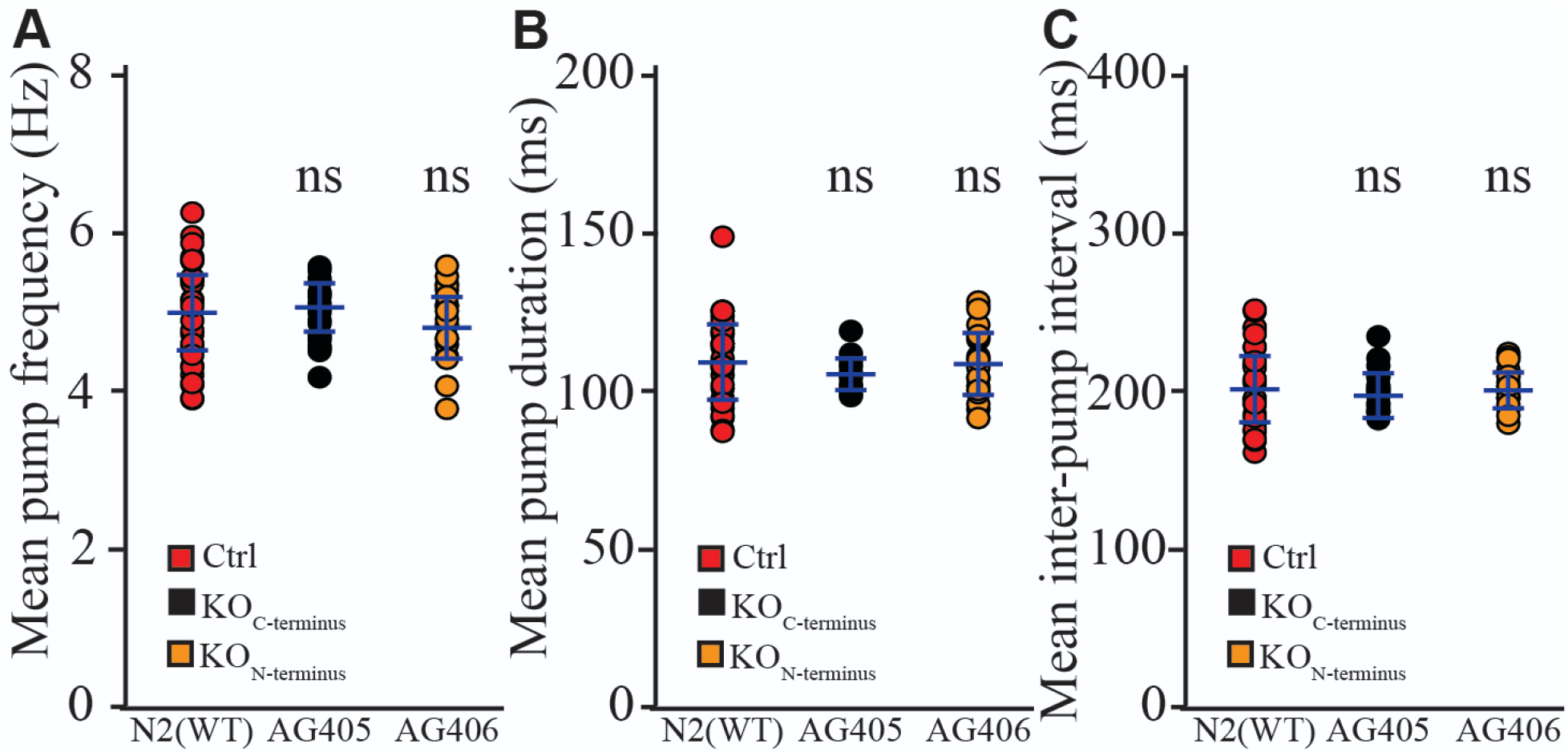
Electropharyngeogram recordings of *pezo-1* mutants induced to perform pharyngeal pumping by immersion in a 10mM serotonin bath displayed no pharyngeal pumping deficits in pumping frequency (**A**), pump duration (**B**), and inter-pump intervals (**C**) N>20. Values reported are means+/− SEMs.

**Supplementary Figure 2.**
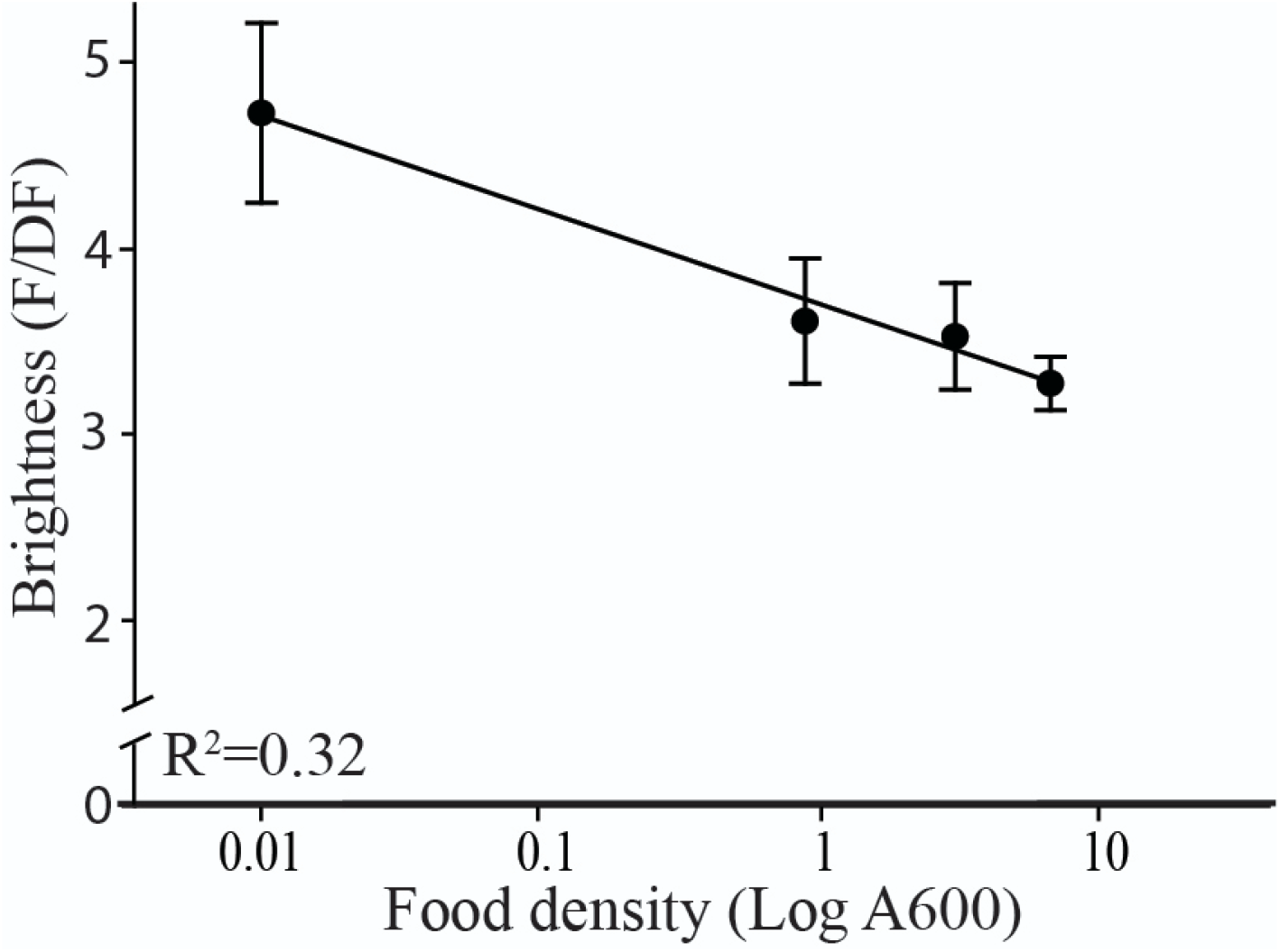
Increasing food density caused the brightness of the pharyngeal gland cells to decrease. NGM agar plates were seeded with a 30 μl lawn of OP50 *E.coli* bacteria at four different densities. Wild type animals (AVG09, N=5-12) were then placed into each plate and allowed to feed for 30 minutes. After this, pharyngeal gland cells were filmed and their brightness measured and normalized against their background. Values reported are means+/− SEMs.

### Supplementary Tables

**Supplementary Table 1.**
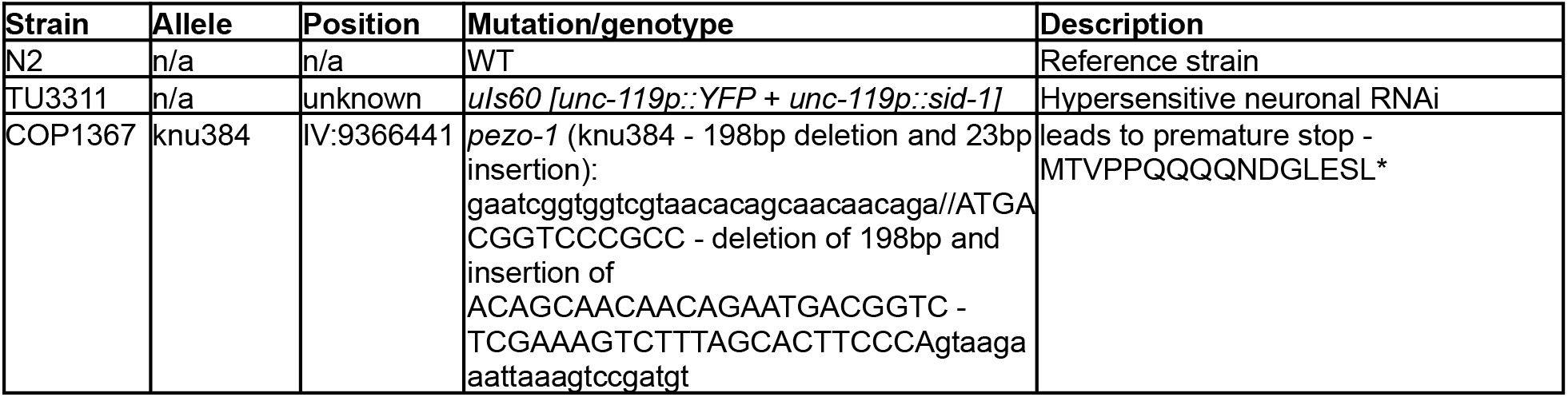
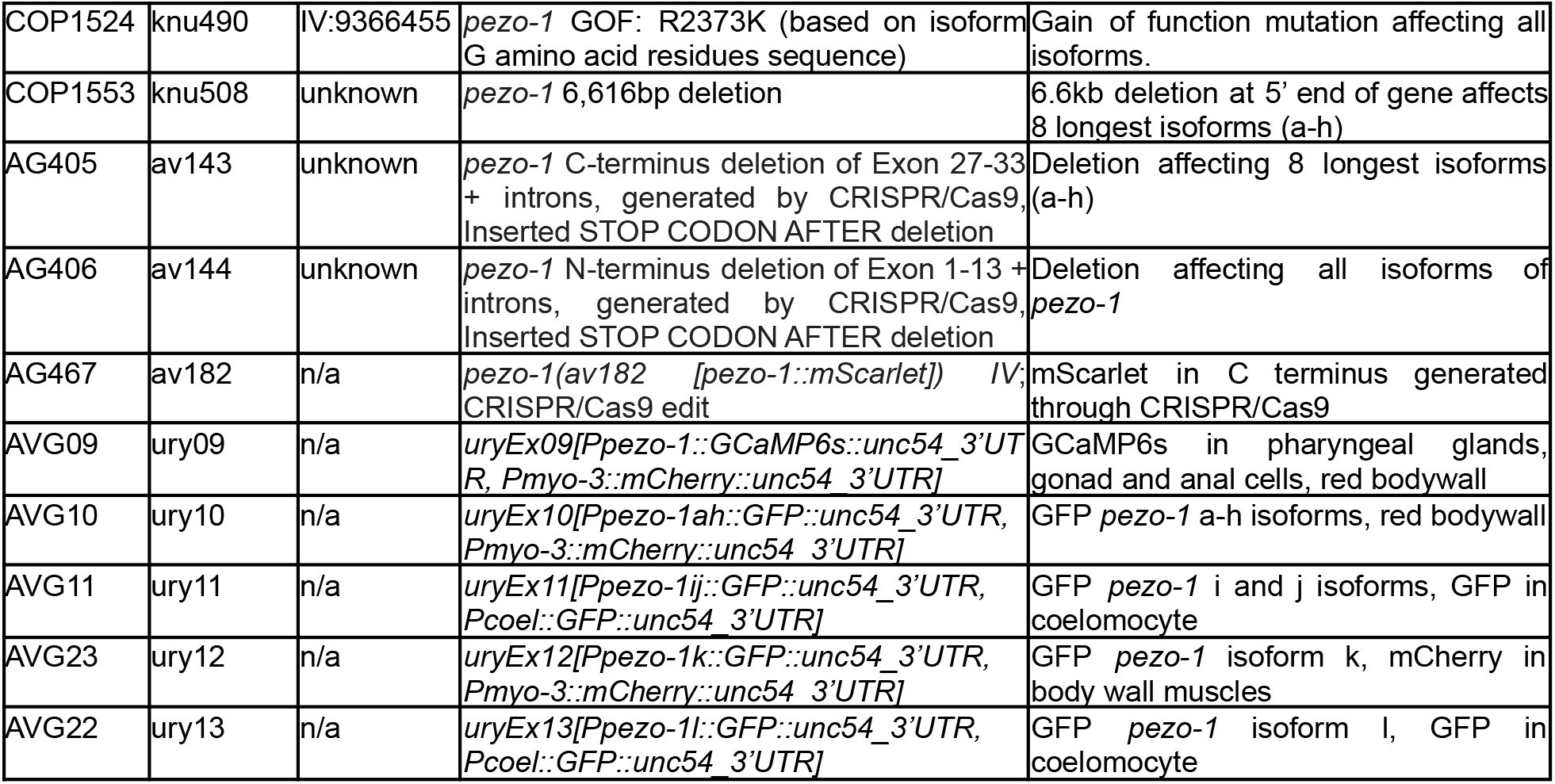
Animal strains used in this study.

**Supplementary Table 2.**
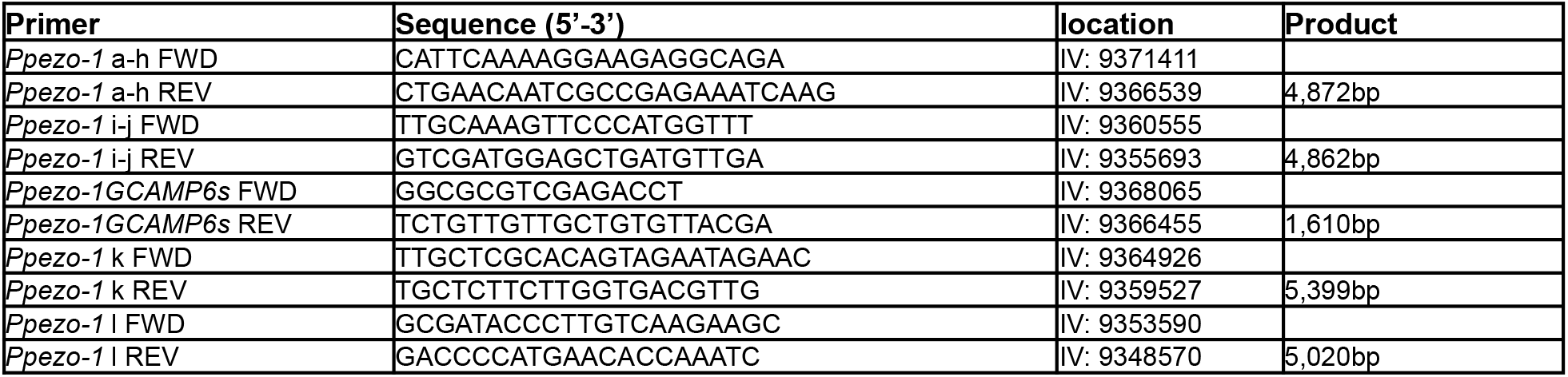
Primers used for the construction of strains used in this study.

**Supplementary Table 3.**
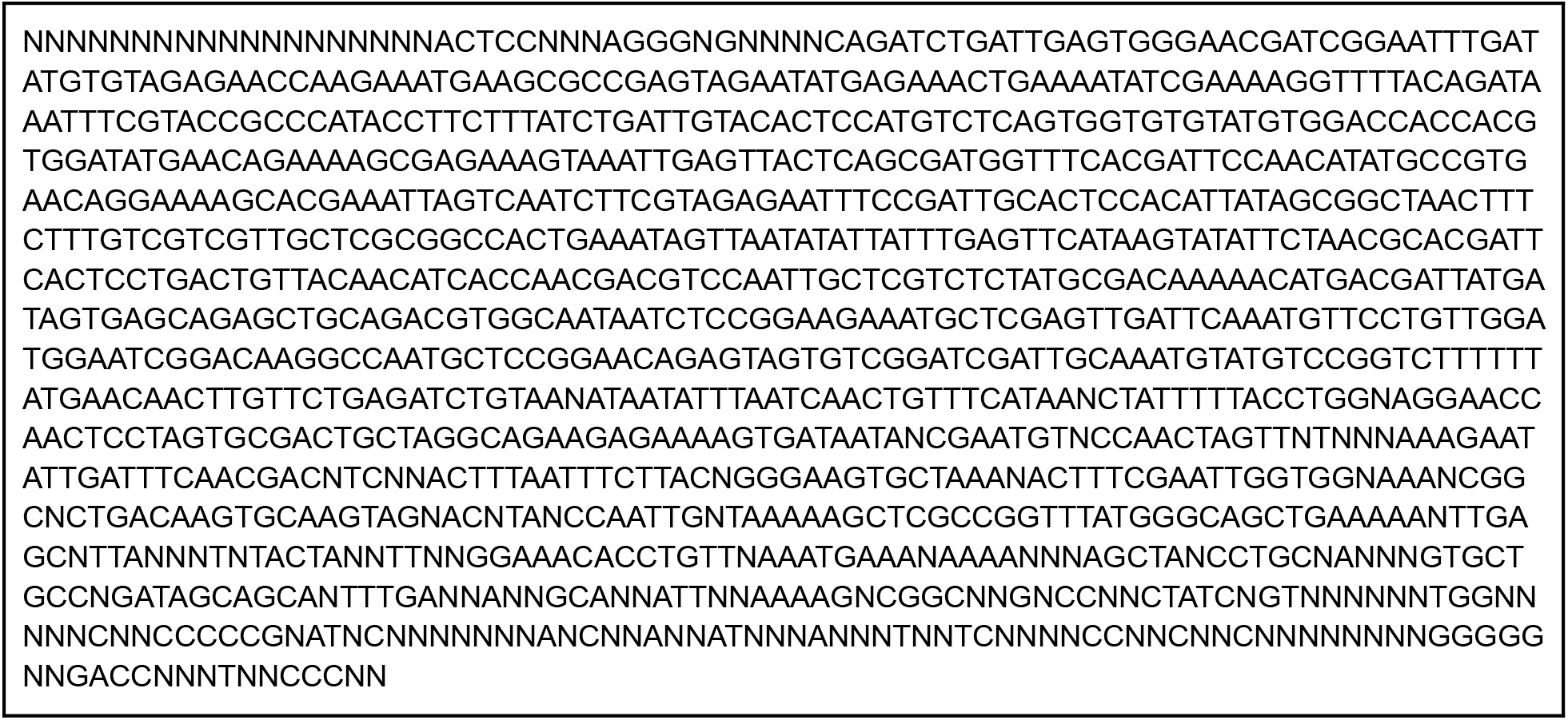
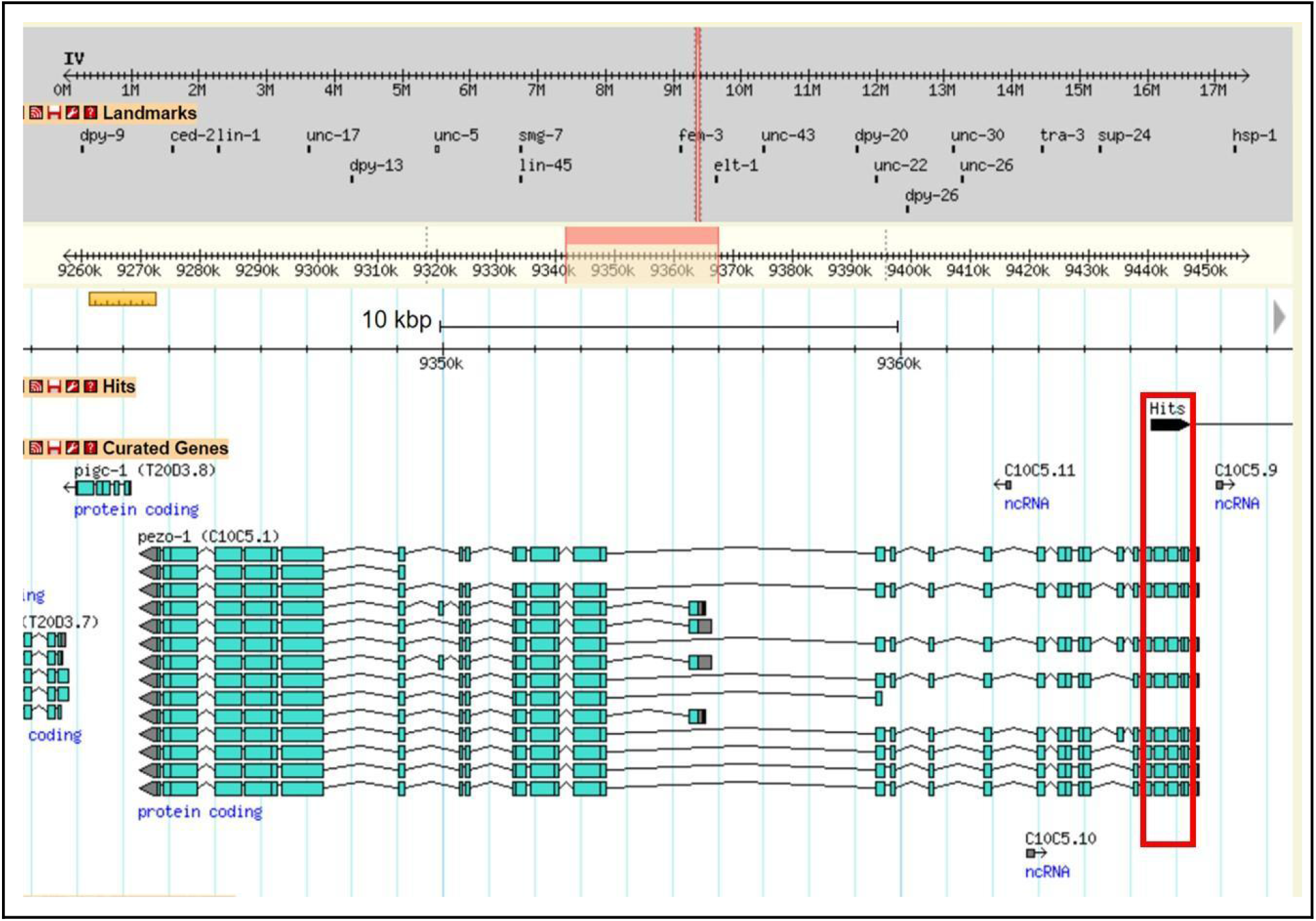
Sequencing results from Source Bioscience for their new *pezo-1* RNAi clone (top), and alignments of the sequence to *C. elegans* genome (bottom) obtained through WormBase through ENSEMBL and BLASTX (Yates et al., 2020; Harris et al., 2020).

**Supplementary Table 4.** Statistical tests used for the different comparisons made in this study. Tests used and sample sizes (for each treatment) for the figures presented in this paper. For Figure 5 and 7D capitalized letters indicate P<0.001 while lower case letters indicate p<0.05. For Figure 6B food density was combined into *low density* (A600=0.32-0.34), *medium density* (A600=2.41-3.49), and *high density* (A600=14.82-15.14).

| Figure | Statistic test | N | Significance |
| --- | --- | --- | --- |
| Figure 5A | One-Way ANOVA with Holm-Sidak test (all pairwise), $F = 16.9$, $DF = 5$ | 10 | *$p < 0.05$ |
| Figure 5B | One-Way ANOVA with Holm-Sidak test (all pairwise), $F = 13.5$, $DF = 5$ | 10-12 | *$p < 0.05$ |
| Figure 6A | Mann-Whitney Rank Sum Test, U statistic = 194 | 29 | **$p < 0.001$ |
| Figure 6B | Two-Way ANOVA (with Holm-Sidak method at significance level = 0.05 ).<br>Strain $F = 13.7$ $DF = 5$ $p < 0.001$; Food density $F = 43.1$ $DF = 2$ $p < 0.001$; Strain x Food density $F = 4.3$ $DF = 10$ $p < 0.001$. Power = 1.0 for three conditions at $\alpha = 0.05$<br>**Food density**<br>N2: High vs Low**, Medium vs Low*<br>AG406: High vs Low**<br>Cop1524: High vs Low**, Medium vs Low*<br>AG405: High vs Low**, High vs Medium*, Medium vs Low**<br>**Strain in high food density**<br>N2 vs (Cop1553**, Cop1367*, Cop1524*)<br>AG406 vs (Cop1553**, Cop1367*)<br>Cop1524 vs (N2*, AG405*)<br>AG405 vs (Cop1553**, Cop1367**, Cop1524*) | 10 | *$p < 0.05$,<br>**$p < 0.001$ |
|  | Cop1367 vs (N2*, AG406**, Cop1524*)<br>Cop1553 vs (N2**, AG405**, AG406**)<br>**Strain in medium food density**<br>N2 vs (Cop1553**, Cop1524**, AG405**, AG406*)<br>**Strain in low food density**<br>Cop1524 vs Cop1367** |  |  |
| Figure 6C | One-Way ANOVA on Ranks (Dunn's on all Pairwise) | 10-26 | *p<0.05 |
| Figure 6D | One-Way ANOVA on Ranks (Dunn's on all Pairwise) | 10-26 | *p<0.05 |
| Figure 6E | One-Way ANOVA on Ranks (Dunn's on all Pairwise) | 10-26 | *p<0.05 |
| Figure 7B | 2nd order regression | >10 | r<sup>2</sup>=0.44 |
| Figure 7C | 2nd order regression | >10 | r<sup>2</sup>=0.23 |
| Figure 7D | Two-Way ANOVA (with Holm-Sidak method at significance level =0.05 ).<br>RNAi treatment F=50.9 DF=1 p<0.001; Food exposure F=11.0 DF=2 p<0.001;<br>RNAi treatment x Food exposure F=13.8 DF=2 p<0.001. Power=1.0 for three<br>conditions at alpha=0.05<br>**Food exposure**<br>RNAi_Ctrl: on food 1 vs (off food**, on food 2*)<br>**RNAi treatment**<br>On food 1: RNAi_Ctrl vs RNAi_pezo-1**<br>On food 2: RNAi_Ctrl vs RNAi_pezo-1* | 10 | *p<0.05,<br>**P<0.001 |
| SuppFig 1A | One-Way ANOVA, F=1.9, DF=2 | >20 | p=0.16 |
| SuppFig 1B | One-Way ANOVA on Ranks | >20 | p=0.24 |
| SuppFig 1C | One-Way ANOVA on Ranks | >20 | p=0.41 |
| SuppFig 2 | 1st order regression | 5-12 | r<sup>2</sup>=0.32 |

**Supplementary Table 5.** Expression profile for *pezo-1* in L1-L4 *C. elegans* larvae obtained through The *C. elegans* Neuronal Gene Expression Map & Network (CeNGEN, Hammarlund et al., 2018). A total of 135 Cells/tissues were found to express *pezo-1* during development. Expression levels were calculated using *n*etwork Differential Gene Expression (nDGE) analysis. Isoforms identified to be expressed in relevant tissues (through PCR-fusion) are indicated.

| Cell | Exp. level | Isoform | Cell | Exp. level | Isoform | Cell | Exp. level | Isoform |
| --- | --- | --- | --- | --- | --- | --- | --- | --- |
| NSN | 338.659 | a-h | RIH | 30.358 |  | RMF | 6.588 |  |
| VA12 | 321.08 |  | PVW | 29.583 |  | SMD stress | 6.424 |  |
| PVD | 273.595 | i-j | AVB | 28.631 |  | RMD DV | 6.091 |  |
| PVT | 233.318 |  | PHA | 28.086 | k | PLM | 5.817 |  |
| CAN | 200.129 |  | AS | 27.483 |  | ASER | 5.685 |  |
| VB | 162.988 | k | AWC OFF | 27.445 |  | FLP | 5.601 | i-j |
| Marginal cell | 112.52 | i-j | AVF | 25.945 |  | Vulva muscle | 5.34 | a-h |
| BDU | 105.389 |  | Glia 2 | 25.194 |  | SIB | 5.301 |  |
| NSM | 105.13 |  | DVC | 25.178 |  | Anal muscle | 5.222 | a-h |
| PVP | 100.069 |  | PVC | 23.789 |  | AIM | 5.13 |  |
| DB | 97.148 |  | Epidermis | 22.811 |  | VB01 | 5.114 |  |
| PHsh | 90.926 |  | ADF | 22.45 |  | CEPsh | 4.859 |  |
| RIR | 86.999 |  | IL2 LR | 21.27 |  | AIB | 4.563 |  |
| PHso | 84.345 |  | CEP | 20.118 |  | AVL | 4.45 |  |
| URA | 77.389 |  | RIC | 19.994 |  | Body wall muscle ant | 3.943 | a-h |
| AMso | 74.719 | a-h | AWC ON | 19.589 |  | VC 4&5 | 3.963 |  |
| VA | 70.921 |  | ASG | 19.085 |  | SMD | 3.803 |  |
| DA9 | 70.243 |  | IL2 DV | 18.652 |  | Unknown cell 2 | 3.63 |  |
| I4 | 69.573 |  | ASK | 18.651 |  | RIS | 3.592 |  |
| VB02 | 68.162 |  | SIA | 18.118 |  | RIB | 3.574 |  |
| Spermatheca | 65.54 |  | Unknown | 17.921 |  | Pharyngeal muscle | 3.412 | a-h |
| Glia 4 | 63.117 |  | ASJ | 17.154 |  | MC | 3.386 |  |
| Glia 1 | 62.07 |  | ADA | 16.9 |  | AFD | 3.376 |  |
| Gon. sheath | 59.384 |  | Germline | 16.375 |  | AWB | 3.355 |  |
| AVD | 57.962 |  | Body wall | 15.647 | a-h, k | ADL | 3.294 |  |
| Glia 3 | 56.962 |  | PDB | 14.718 |  | URX | 3.113 |  |
| DA | 54.272 |  | VD DD | 14.571 |  | AIN | 3.05 |  |
| Arcade cell | 53.602 | k | Coelom | 14.14 |  | RMD LR | 2.907 |  |
| DB01 | 52.126 |  | Rectal gland | 13.392 | a-h | BAG | 2.791 |  |
| RIP | 45.997 |  | AVE | 12.256 |  | SAB | 2.334 |  |
| Sperm | 45.68 |  | AVJ | 12 |  | VC | 2.178 |  |
| DVB | 45.997 |  | ASEL | 11.957 |  | AUA | 2.03 |  |
| M2 | 43.768 |  | Spermatheca | 11.805 |  | AMsh | 1.987 | a-h |
| Unkn. cell 1 | 43.121 |  | ASH | 11.654 |  | AVK | 1.787 |  |
| Dist. tip cell | 41.971 |  | PLN | 11.441 |  | AIY | 1.7 |  |
| Unannotated | 39.01 |  | RMH | 11.336 |  | RIA | 1.51 |  |
| AIA | 37.666 |  | Excretory | 10.749 | i-j | Vulval cells | 1.364 |  |
| Uterine | 36.89 | a-h | Seam cell | 10.43 |  | RIM | 1.333 |  |
| Intestine | 35.362 | i-j | RIV stress | 10.392 |  | AVM | 1.253 |  |
| ASI | 34.146 |  | Pharyngeal gland | 10.006 | a-h | PVQ | 1.227 |  |
| AVG | 33.602 |  | AVA | 9.402 |  | RIV | 1.156 |  |
| I3 | 33.109 | a-h, i-j | AVH | 8.986 |  | RID | 1.056 |  |
| LUA | 31.109 |  | SMB | 8.698 |  | M5 | 0.749 |  |
| PHB | 30.693 |  | ALM | 8.441 |  | RIG | 0.549 |  |
| AWA | 30.638 |  | l1 | 7.236 |  | AIZ | 0.539 |  |

